# Poly-L-Ornithine Coated Plant Scaffolds Support Motor Recovery in Rats after Traumatic Spinal Cord Injury

**DOI:** 10.1101/2025.02.05.636658

**Authors:** Lauren J. Couvrette, Krystal L. A. Walker, Arman Bayat, Daniel J. Modulevsky, Alex M. Laliberté, Charles M. Cuerrier, Maxime Leblanc-Latour, Ryan J. Hickey, Rose Boudria, Ruthie Monty, Ras-Jeevan K. Obhi, Isabel Shore, Ahmad Galuta, Eve C. Tsai, Tuan V. Bui, Andrew E. Pelling

## Abstract

Spinal cord injury (SCI) is a debilitating neurological condition with far-reaching consequences for patients, including loss of motor function and significant limitations to quality of life. Implantable biomaterials have emerged as a therapeutic strategy to modulate the SCI microenvironment and facilitate regeneration of axons. In this study, plant-derived lignocellulosic scaffolds coated with poly-L-ornithine (PLO) are shown to support locomotor recovery and neural tissue repair in a rat model of spinal cord injury. Upon complete transection of the spinal cord, animals were implanted with a plant-derived scaffold coated in poly-L-ornithine, a positively charged amino acid chain that is known to promote neural stem cell differentiation into neurons and enhance myelin regeneration. Recovery of motor function was evaluated by the Basso, Beattie and Bresnahan (BBB) locomotor scale as well as the Karolinska Institutet Swim Assessment Tool (KSAT). Retrograde tracing of ascending sensory tracts revealed enhanced regeneration in animals that received the PLO-coated scaffold. Numerous β-III tubulin and neurofilament 200 positive fibers may indicate axonal sprouting within the lignocellulosic scaffold and LFB staining highlights myelination around the PLO-coated scaffold. These results demonstrate the potential of plant-based biomaterials in a rat model of acute spinal cord injury and highlight their enhancement after PLO functionalization.

## Introduction

The mammalian spinal cord has limited regenerative capacity after spinal cord injury (SCI). Failure of the adult central nervous system to regenerate can be attributed in part to the limited capacity of injured neurons to re-establish functional axons across the lesion^1^. Mechanical trauma to the tissue elicits a range of cell responses, including increased proliferation of ependymal cells that line the central canal^2^. Upon activation, ependymal cells re-express neural stem cell properties and migrate to the site of injury, where they spontaneously differentiate into oligodendrocytes and astrocytes^3^. Although ependymal cells can differentiate into neurons *in vitro*, the same has not been demonstrated following SCI in vivo^4^, though mature oligodendrocytes have been observed^5,6^. After SCI, complex pathological mechanisms create a hostile microenvironment at the injury epicenter, which inhibits the regrowth of axons and damages surrounding tissue^7^. For instance, SCI-induced death of oligodendrocytes leads to an accumulation of myelin breakdown products at the injury site ^8–10^. Molecules such as Nogo-A and myelin-associated glycoprotein are released from damaged myelin into the extracellular matrix after SCI and represent an important group of axonal growth inhibitors^11^. Many therapeutic strategies aim to establish a more permissive environment for regeneration by removing inhibitory molecules^12,13^, providing trophic support^14,15^, directing stem cell fate^16^, remyelinating axons^17,18^, or implanting biomaterial scaffolds that can guide axonal growth^19,20^. In recent years, multimodal therapies have generated much attention since they address multiple aspects of SCI pathology and often produce greater benefit than their individual components^21,22^. Such therapeutic strategies have combined cell transplants, biomaterials, locomotor training, and neurotrophic factors with the goal of modulating the injury microenvironment to promote repair.

Developing biocompatible scaffolds to bridge the SCI has been at the forefront of tissue engineering strategies. Implantable biomaterials not only provide structural scaffolding to guide cell attachment and migration but can also be used to regulate the inflammatory response or deliver other therapeutics such as stem cells and growth factors. Regenerative biomaterials for SCI are designed to replicate the properties of spinal cord tissue and should therefore have the mechanical strength, porosity and internal microstructures that are suitable for neurite extension across the injury site^23,24^. Moreover, the surface chemistry of biomaterials is of particular importance, as it can be utilized to influence cell migration and differentiation ^25,26^.

Both degradable and non-degradable biomaterials have been investigated for SCI repair. In order for degradable scaffolds to act as extracellular matrix substitutes, it is essential that their rate of degradation aligns with the regeneration rate of axons^27^. Synthetic materials are attractive candidates for spinal cord tissue engineering due to their controllable mechanical properties and high batch-to-batch consistency. For example, polyethylene glycol is a synthetic polymer that can be cross-linked to form injectable hydrogels that provide a framework for regenerating tissue and were shown to promote motor recovery in a rat model of SCI^28^. In addition, poly(lactic-co-glycolic acid)-based scaffolds have produced significant motor improvements and increased tissue remodelling in African green monkeys with an incomplete SCI^29^. However, many synthetic biomaterials are limited by their hydrophobicity, which can impair cell attachment, or by their potential to release toxic by-products during degradation. By contrast, natural materials such as collagen, fibrin, alginate, chitosan and hyaluronic acid are widely studied as regenerative scaffolds for SCI due to their excellent biocompatibility and cell adhesion properties. In particular, collagen-based scaffolds are a promising SCI treatment that has been explored in diverse forms including collagen hydrogels^30^, sponges or scaffolds that deliver therapeutics to the injury^31^. Ongoing human clinical trials have reaffirmed the therapeutic potential of collagen scaffolds, notably in their ability to promote tissue regeneration and functional recovery^32^. One common issue with scaffolds made from natural polymers is insufficient mechanical strength and durability in vivo, which multiple studies have aimed to resolve by adding cross-linkers^33^ or combining natural and synthetic biomaterials.

Recently, plant-derived scaffolds^34–41^ have been investigated in vivo for various biomedical applications including skin^42^, tendon^43^, nerve^44^ and bone^45^ tissue engineering. Many different plant species can be decellularized with the help of detergents, sonication, enzymatic digestion, or freeze-thawing methods to produce cellulose-based scaffolds^46^ composed of β(1→4) linked D-glucose units. Plant cellulose scaffolds were shown to be biocompatible in vivo and supported extracellular matrix deposition when implanted subcutaneously^34^. In addition, these scaffolds became vascularized as early as one-week post-implantation. Since cellulose is not biodegradable in humans, scaffolds made from this material are offer long-term durability in vivo^35^. Moreover, plant cellulose has proven to be a versatile material that can be combined with hydrogels to generate customizable macroscopic structures^36^. Cellulose scaffolds can also be functionalized with various coatings to promote cell adhesion^47^. For example, we have previously demonstrated that PLO-coated lignocellulosic scaffolds produced from stalks of Asparagus officinalis can support attachment and proliferation of neural stem cells in vitro^37^. In culture, rat neural stem cells were shown to form neurospheres that attach to asparagus-derived scaffolds and readily infiltrate into its vascular architectures. Increased expression of neuron-specific beta-III tubulin and glial fibrillary acidic protein in cells grown on these scaffolds suggests they may promote differentiation of NSCs towards neurons and astrocytes, respectively, *in vitro*.

Here we investigate the ability of PLO-coated lignocellulosic biomaterials to support neural tissue repair and functional recovery after traumatic SCI. We implanted PLO-coated scaffolds or uncoated scaffolds acutely following a complete spinal cord transection in rats housed in an enriched environment. Motor recovery was monitored by the Basso, Beattie and Bresnahan (BBB) open field assessment and Karolinska Institutet Swim Assessment Tool (KSAT). Sprouting and regeneration of the dorsal columns were analyzed by neural tract tracing with the retrograde tracer CTb injected into the sciatic nerve. The results of this experiment point to enhanced regeneration of the sensory tracts in animals that received PLO-coated implants compared to controls. In addition, the descending fibers of the corticospinal tracts were visualized with dextran amine injections in the motor cortex, which revealed CST axons projecting along the lignocellulosic biomaterial. Finally, histological analysis showed host tissue integration with the scaffold and neural cell migration along its channels, which was enhanced by the PLO coating. In sum, we demonstrate the ability of PLO-modified plant-derived scaffolds to promote axonal sprouting in the lesion and induce motor recovery after SCI.

## Results

### Lignocellulosic Scaffold Production and Implantation in Rodent Spinal Cord Injury

Stalks of Asparagus officinalis were cut into cylindrical sections with a biopsy punch and decellularized to produce highly structured three-dimensional lignocellulosic scaffolds as confirmed through imaging (Fig 1A) and FTIR analyses (Supplementary Information Fig 1 and Supplementary Information Table 1). Characterization of the plant-derived scaffolds are briefly described here but were also reported in a prior study^37^. Each scaffold contains naturally occurring vascular bundles (VBs) (Fig 1B, C), which are circularly arranged channel-like structures spaced 612±70μm (n=11) apart by parenchyma tissue. The parenchyma tissue is highly porous with an average pore size of 39±15µm (n=8) (Supplementary Information Fig 2). The VBs are composed of characteristic channel-like structures known as xylem, sieve tubes and phloem which have average diameters of 51±15 μm (n=11), 40±16μm (n=11) and 9±2μm (n=11) respectively. Each prepared scaffold had 11±2 VBs (n=4) which contain 35±5 continuous microchannels (n=11) (Supplementary Information Fig 2) that linearly span the entire scaffold (Supplementary Information Fig 3). The decellularized scaffolds were mechanically characterized and found to be anisotropic due to the linear and highly aligned architecture of the VBs which run parallel to the plant stem. When measured parallel or perpendicular to the plant stem, the elastic modulus was found to be 148±53kPa (n=10) or 12±4 kPa (n=10) (Supplementary Information Fig 4). The mechanical properties of the scaffolds utilized in this study are broadly consistent with reported values for the spinal cord although these values can be highly variable and dependent on numerous factors^48–52^.

**Figure 1.**
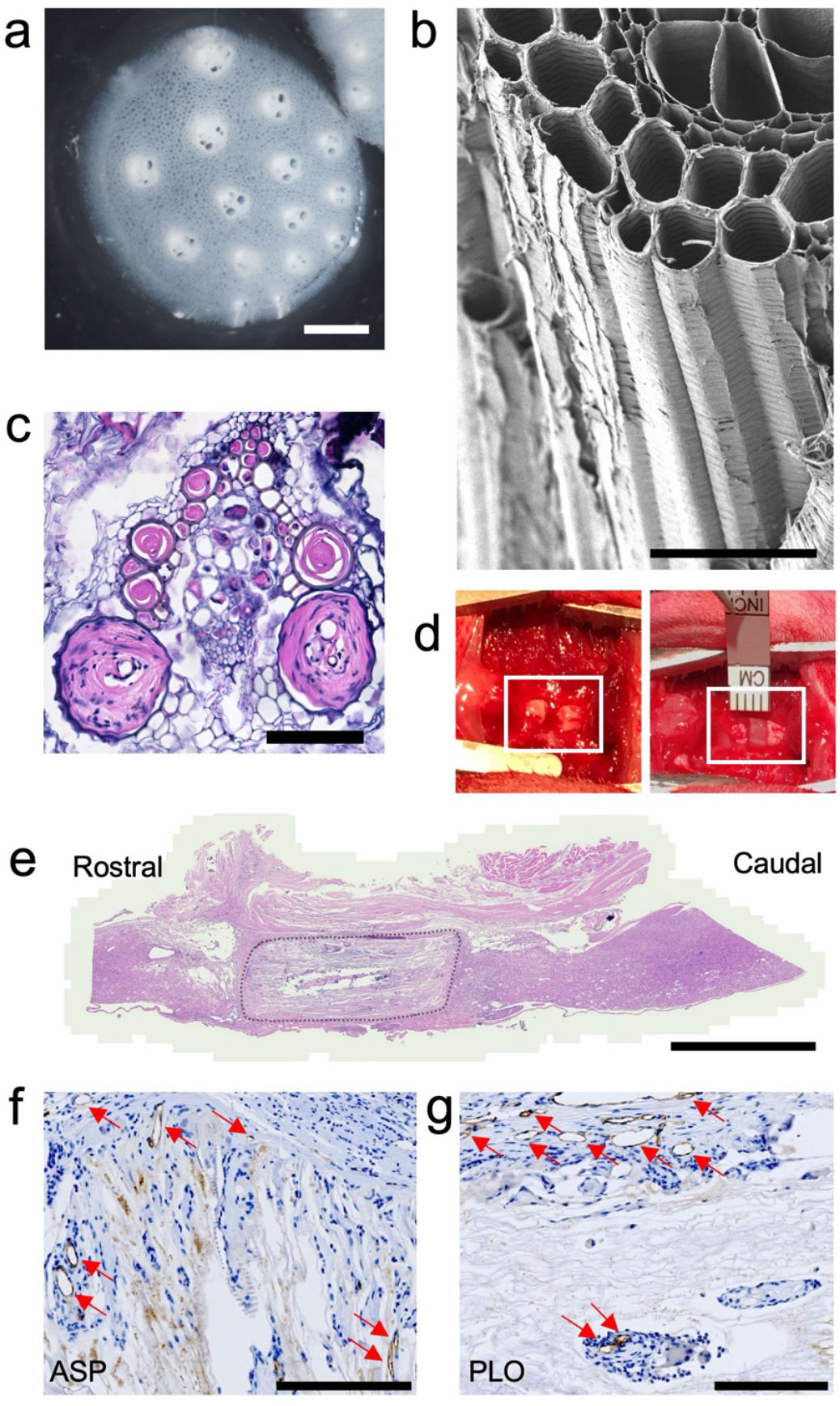
Plant-derived lignocellulosic biomaterial in rodent model of complete transection spinal cord injury. A. A cross sectional view of decellularized plant-derived scaffold consisting of vascular bundles and parenchyma (scale bar = 1 mm). B. SEM of lignocellulosic scaffold microarchitecture (scale bar = 100μm). C. Hematoxylin & eosin staining of cross section of lignocellulosic scaffold after 12 weeks in vivo. The scaffold’s vascular bundles are infiltrated with host cells. (Scale bar = 200 µm). D. An exposed fully transected spinal cord (left box) and a scaffold after implantation (right box). E. Hematoxylin & eosin staining of sagittal section of spinal cord tissue with lignocellulosic scaffold (outlined) after 12 weeks in vivo (Scale bar = 2mm). F. Immunohistochemistry staining for CD31 (brown) in a sagittal section at the caudal biomaterial-tissue interface (representative image from the ASP group). CD31 positive blood vessels denoted by arrow (Scale bar = 200µm). G. Immunohistochemistry staining for CD31 (brown) in a sagittal section at the caudal biomaterial-tissue interface (representative image from the PLO group). CD31 positive blood vessels denoted by arrow (Scale bar = 200µm).

In animals with a complete T8-T9 spinal cord transection, scaffolds were implanted into the gap between the stumps of the cord with the long axis of the VBs parallel to the spinal cord (Fig 1D). We utilized scaffolds with a diameter of 4 mm and an average length of 2.9±0.5mm which was customized to fit the specific dimensions of each animal’s injury. The control group (n=8) did not receive a scaffold, while the PLO group (n=11) was implanted with a lignocellulosic scaffold coated with a 100µg/ml solution of PLO and uncoated scaffolds were implanted into the ASP group (n=9). After 12 weeks in vivo, spinal cord tissue was collected, and histological analysis was performed to assess tissue integration with the scaffold. Hematoxylin & eosin staining revealed nuclei migrating along the channels of the biomaterial (Fig 1C, E), which is consistent with our previous in vitro study^37^. In particular, nuclei were observed within the vascular bundles of the biomaterial (Fig 1C) and across the lesion. Immunohistochemistry staining for CD31 was performed to assess biomaterial vascularization (Fig 1F, G). CD31 positive blood vessels were identified in the tissue surrounding the scaffold, including the dorsal, rostral, and caudal interfaces in both the PLO and ASP groups. As well, CD31 positive blood vessels were found inside the vascular bundles within the biomaterial. The scaffolds retained their structure and dimensions throughout the study, and no signs of chronic foreign body reaction were observed, which aligns with the findings of previous studies^34,53^. Moreover, prior to testing the biomaterial in this SCI model, we also examined its biocompatibility by subcutaneously implanting PLO-coated scaffolds in immunocompetent Sprague Dawley rats. Histological analysis performed at 1-, 4-, and 12-weeks post-implantation showed a gradual decrease in immune cell presence in the scaffold and surrounding dermis tissue (Supplementary Information Figure 5). By the 12-week timepoint, only very low levels of foreign body multinucleated cells were observed at the periphery of the scaffold.

### Hindlimb locomotor recovery after complete SCI

#### BBB Locomotor Assessment

Hindlimb motor function was assessed weekly using the Basso, Beattie and Bresnahan (BBB) locomotor scale, which ranges from 0 (complete hindlimb paralysis) to 21 (normal locomotion) points. Two weeks after complete transection of the spinal cord, the BBB assessment was performed to confirm total hindlimb paralysis. At this initial assessment, there were no significant differences in locomotor ability amongst the experimental groups: No Tx (transection with no scaffold, n=8), ASP treated (lignocellulosic implant only, n=9), and PLO treated (PLO-coated lignocellulosic implant, n= 11). Any animal that achieved a BBB score of above 5 at the 2-week post-operative timepoint was excluded from further locomotor evaluation. Out of 29 animals, a single rat was identified as an outlier, with a BBB score of 5.5 (average of 3 examiners) and was therefore excluded from the BBB motor recovery assessment. Over the 11-week recovery period, BBB scores increased significantly in animals that received PLO-coated scaffolds (p<0.0001) as well as those implanted with uncoated scaffolds (p=0.0158) (Fig 2A). This was determined by comparing the average score at week 2 with the average score at week 11 for each experimental group. Whereas significant functional recovery was achieved in scaffold treated groups, animals that did not receive a scaffold had no significant change (p=0.3128) in BBB scores throughout the recovery period. At week 11, the mean BBB score of rats treated with the PLO-coated scaffold was 5.6±0.87 points, corresponding to slight movement in 2 joints and extensive movement of a third. The group receiving uncoated scaffolds had a mean score of 4.3±0.96, indicating slight movement of all three joints of the hindlimb. By contrast, controls that did not receive a scaffold had a mean score of 3.35±1.0, reflecting extensive movement in 2 joints. Importantly, no animals from the No Tx group attained scores of 6 or above, whereas several animals from the PLO and ASP groups achieved BBB scores of 7 (extensive movement of all three joints), 8 (plantar placement of the paw without weight support), and 9 (plantar placement of the paw with weight support in stance only).

**Figure 2.**
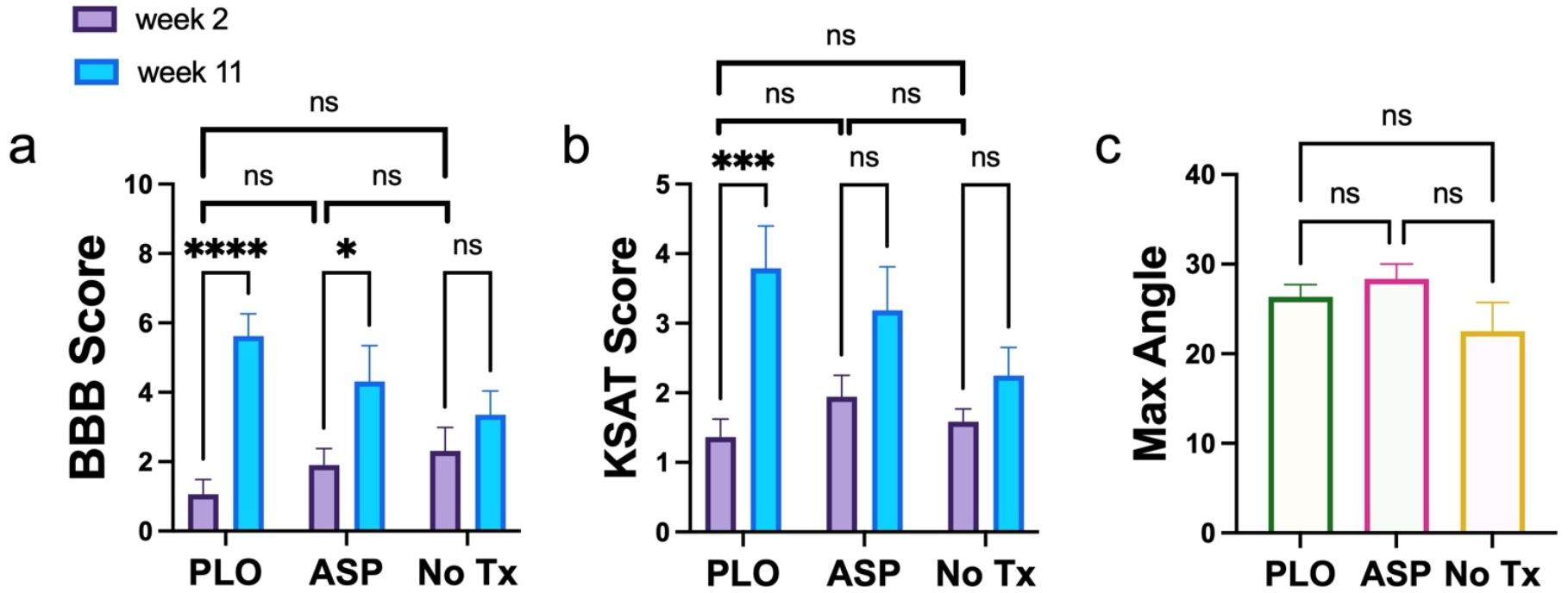
Locomotor assessments after complete spinal cord transection. A. Average BBB score of each experimental group at week 2 after SCI compared to week 11 after SCI (2way ANOVA Uncorrected Fisher’s LSD; ****P<0.0001, *P=0.0158; mean ± s.e.m, *n* = 11 PLO animals, n = 9 ASP animals, n = 8 No Tx animals). B. Average KSAT swim assessment score of each experimental group at week 2 after SCI compared to week 11 after SCI (2way ANOVA Uncorrected Fisher’s LSD; ***P=0.0001; mean ± s.e.m, *n* = 11 PLO animals, n = 9 ASP animals, n = 8 No Tx animals). C. Average maximum angle achieved by each experimental group in inclined plane assessment at final week (mean ± s.e.m, n = 11 PLO animals, n = 9 ASP animals, n = 8 No Tx animals).

#### KSAT Swimming Assessment

Motor recovery as assessed by the BBB can be influenced by the retraining effect^54–56^, whereby spontaneous “self-training” occurring in the cage contributes to functional recovery. Factors such as cage size and housing density can influence the activity level of the animals, which may impact the recovery of overground walking. By contrast, swimming performance is not affected by the retraining effect^57^, since animals are not normally exposed to swimming in a laboratory setting. Another potential confounding factor that may affect BBB scores is sensory feedback from below the injury, which can contribute to motor recovery during overground walking after SCI^58–61^. During swimming, there is an important reduction in sensory feedback from the hindlimbs compared to overground walking^57,62^ since rats rely on buoyancy to support their bodyweight. The Karolinska Institutet Swim Assessment Tool (KSAT) evaluates swimming parameters including hindlimb movement, forelimb usage, trunk instability, and body angle. In this assessment, healthy animals achieve a maximum score of 19, based on intensity and frequency of limb and tail movement. At the first post-operative swim assessment, there were no differences amongst the experimental groups. After 11 weeks of recovery, rats with the PLO-coated scaffold showed significant improvements in swimming performance compared to their first post-operative assessment (p=0.0001) (Fig 2B). By contrast, the animals treated with uncoated scaffolds (p=0.0609) and those without scaffolds (p=0.3361) did not achieve statistically significant improvement over the course of the study. At the final swimming assessment, the average KSAT score of rats implanted with PLO-coated scaffolds was 3.79± 0.58 compared to 3.18±0.64 with the uncoated scaffold (ASP group) and 2.25±0.68 for those without scaffolds (No Tx group). Overall, animals from the PLO and ASP groups showed improvement in hindlimb movement and trunk stability. Interestingly, the best performing animal from the PLO group attained a KSAT score of 8, while the highest score achieved in the ASP group was 6 and the highest score in the No Tx group was 5.

#### Inclined Plane test

Before the spinal cord injury, all animals were trained to perform the inclined plane test as described by Rivlin et al^63^. Animals were placed on an inclined plane and the slope was adjusted to determine the maximum angle at which the animal could maintain its position without falling. Every 2 weeks after the spinal cord injury, the inclined plane test was performed to examine sensorimotor recovery. At the final week, there were no significant differences (p= 0.7915) in the performance of the three experimental groups (Fig 2C). On average, animals reached a maximum angle of 26±1 degrees in the PLO group (PLO-coated scaffolds), 28±1 degrees in the ASP group (uncoated scaffolds), and 23±3 degrees in the control group (No scaffold).

### Retrograde tract tracing of ascending sensory fibers

To elucidate possible mechanisms that may facilitate the observed motor recovery, retrograde tracing was performed on rats from each experimental group. Neuroanatomical tract tracing is commonly used to visualize sprouting or regeneration of axons after injury^64–67^. After the recovery period of 11 weeks, Cholera Toxin subunit B (CTb) conjugated to Alexa Fluor 647 was injected into the sciatic nerve to label sensory axons and measure their projection distance along the dorsal column of the spinal cord (Fig. 3A). Cross sections of T10 spinal cord tissue caudal to the injury were imaged by confocal laser scanning microscopy and revealed a distinct CTb-positive signal in the dorsal column (Fig 3B) which was not found in T6 sections rostral to the injury. Sagittal sections of spinal cord tissue showed CTb-labelled axons projecting into the caudal side of the scaffold towards the injury epicenter (Fig 3C-E). Compared to animals without a scaffold, those with PLO-coated scaffolds had on average a significantly smaller distance (p=0.0264) between the furthest rostral CTb-traced axons and the injury epicenter (Fig 3A). This result suggests that the PLO-coated scaffold enhanced regeneration of axons and/or reduced axon retraction. In animals treated with PLO-coated scaffolds (Fig 3E), the furthest rostral CTb-labelled axons were identified at an average of 3.2±0.2 mm from the injury epicenter. By contrast, control animals that did not receive a scaffold had CTb-traced axons at an average distance of 4.8±0.4 mm from the epicenter (Fig 3C). In animals treated with the uncoated scaffold, the average distance of CTb-traced axons from the epicenter was 3.2±0.3 mm. This tract tracing experiment reveals that sensory axons extend further rostrally in animals with PLO-coated scaffolds compared to those without a scaffold. However, it is unclear whether those sensory fibers are regenerated axons or spared fibers that may result from reduced cystic cavitation after injury.

**Figure 3.**
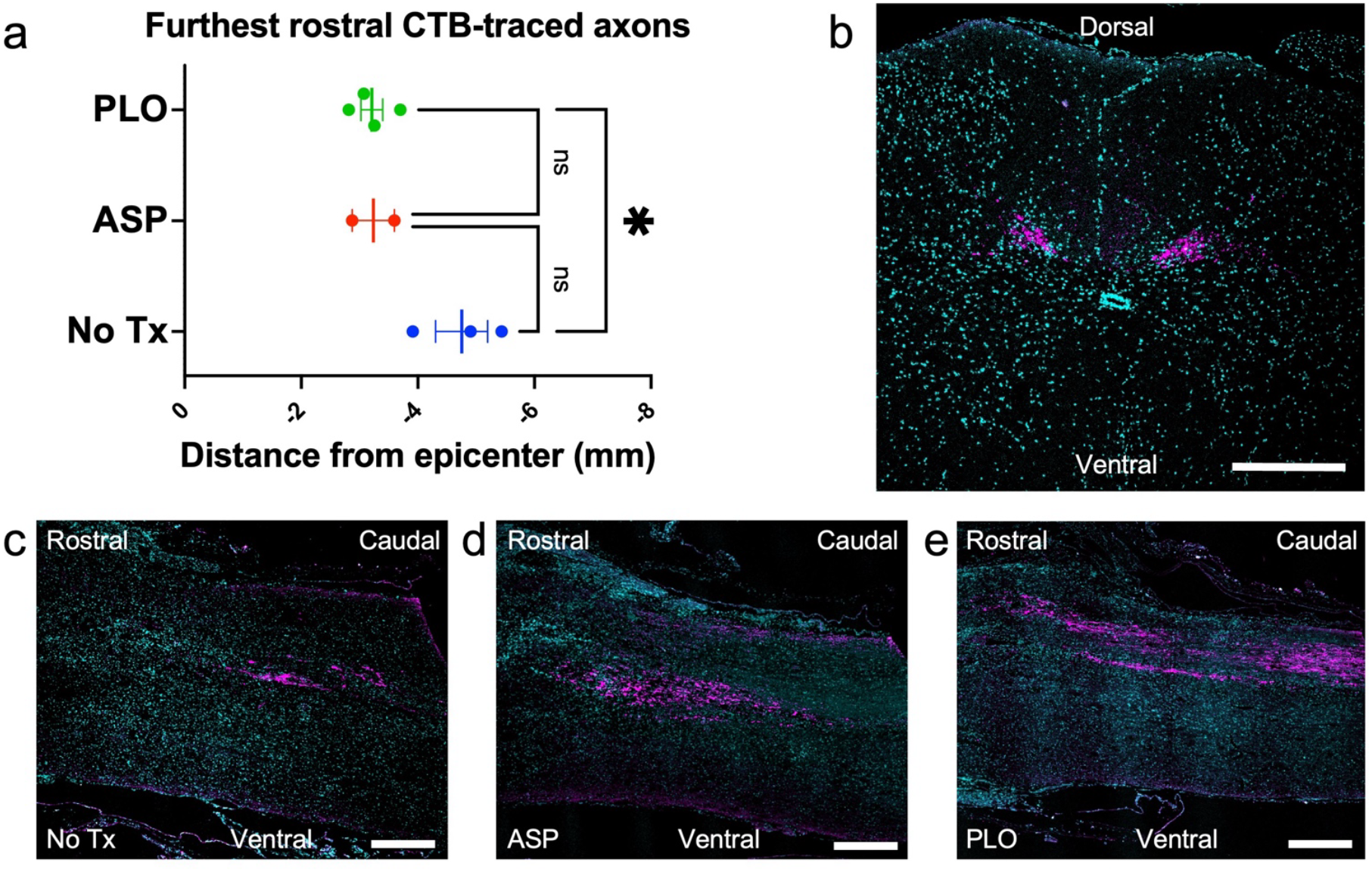
Retrograde tract tracing of ascending sensory afferents by CTb injection into sciatic nerve. A. Furthest rostral CTb-traced axons. Each point represents the distance (mm) between the injury epicenter and the furthest rostral CTB-traced axon in one animal. Bar represents the mean ± s.e.m of each group (One-way ANOVA; *P=0.0264, n = 4 PLO animals, n = 2 ASP animals, n = 3 No Tx animals). B. Hoescht (cyan) stained cross section of spinal cord at T10 (caudal to the injury) confirms presence of CTb (magenta) in dorsal column and lamina 4 (Scale bar 300 µm). C. Hoescht (cyan) stained sagittal section of spinal cord from SCI animal with no scaffold at T9 (caudal side of the injury). CTb-traced axons (magenta) can be seen in the dorsal collumn (Scale bar 500 µm). D. Hoescht (cyan) stained sagittal section of spinal cord from SCI animal with uncoated scaffold at T9 (caudal side of the injury). CTb-traced axons (magenta) can be seen in the dorsal collumn (Scale bar 500 µm). E. Hoescht (cyan) stained sagittal section of spinal cord from SCI animal implanted with PLO-coated lignocellulosic scaffold at T9 (caudal side of the injury). CTb-traced axons (magenta) can be seen in the dorsal column (Scale bar 500 µm).

### Anterograde labeling of the corticospinal tract

Another tract tracing experiment was performed to investigate the regeneration of the hindlimb corticospinal tract (CST), which is involved in the control of voluntary movement in mammals^68,69^. After the recovery period of 11 weeks, stereotaxic injections of dextran amine (DA) conjugated to Alexa Fluor 488 were made into the hindlimb motor cortex, to measure the distance of descending projections in the spinal cord (Fig. 4A). Successful tracer uptake was confirmed by confocal laser scanning microscopy of T6 spinal cord cross-sections rostral to the injury, which showed strong positive dextran amine labelling of axons in the corticospinal tract (Fig 4B). In sagittal sections, dextran amine-labeled axons were identified at the rostral interface between the biomaterial and spinal cord (Fig 4D & E), though none were present in T10 cord sections caudal to the injury. The most caudal DA-labelled axons were identified at an average distance of 0.33±0.4 mm from the epicenter for animals with PLO-coated scaffolds (Fig 4E), 0.45±0.3 mm from the epicenter for those with the uncoated scaffold (Fig 4D), and 1.6±0.1 mm from the epicenter for animals without a scaffold (Fig 4C). Although no significant difference in CST axon extension was found between the experimental groups, DA-traced axons were seen projecting along the dorsal side of the PLO-coated lignocellulosic biomaterial towards the caudal spinal cord (Fig 4E).

**Figure 4.**
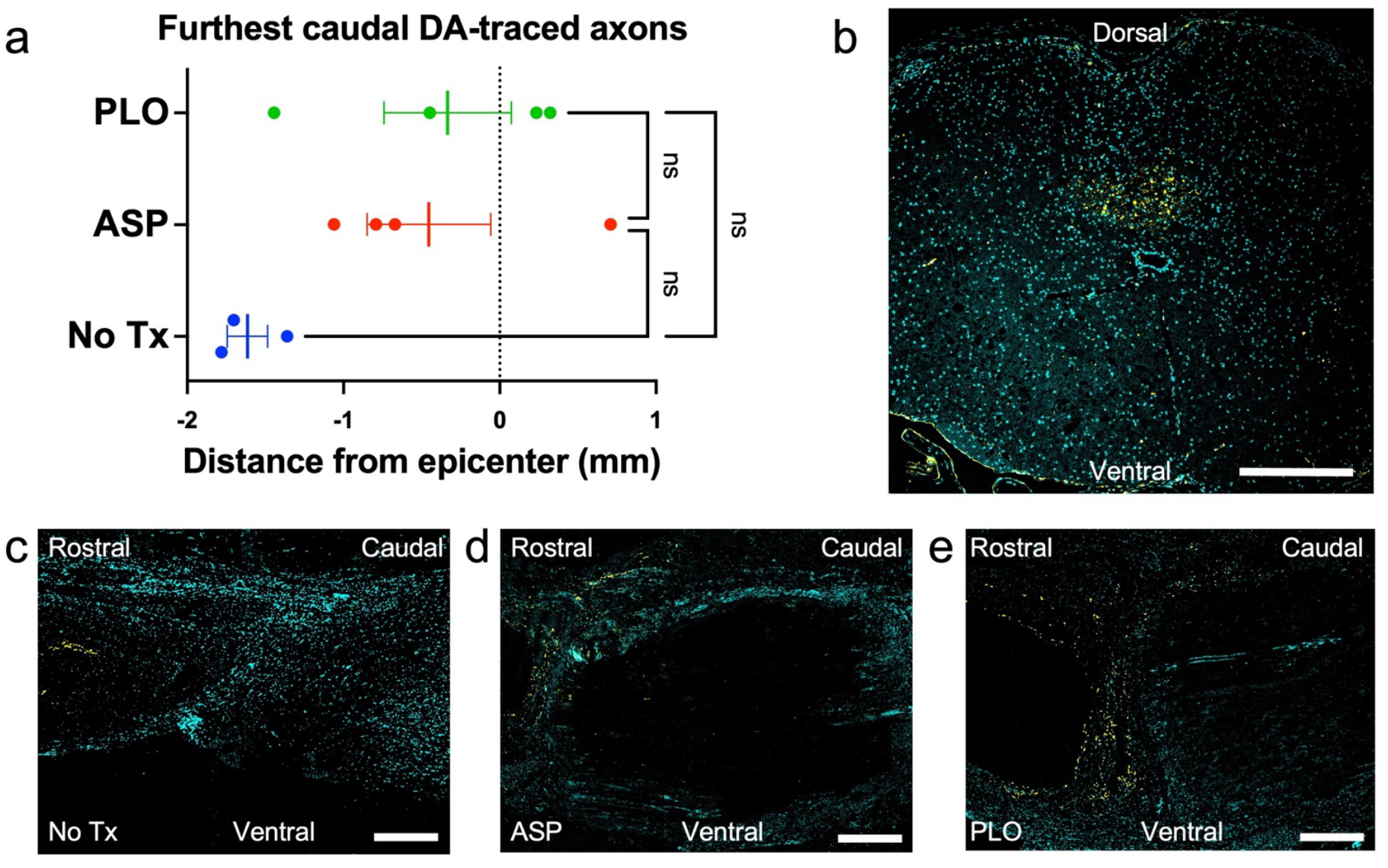
Anterograde labeling of the corticospinal tract by injection of dextran amine into hindlimb motor cortex. A. Furthest caudal dextran amine-traced axons. Each point represents the distance (mm) between the injury epicenter and the furthest caudal DA-traced axon in one animal. Bar represents the mean ± s.e.m of each group (One-way ANOVA; P=0.0903, n = 4 PLO animals, n = 4 ASP animals, n = 3 No Tx animals). B. Hoescht (cyan) stained cross section of spinal cord at T6 (rostral to the injury) confirms presence of dextran amine (yellow) in corticospinal tract (Scale bar 300 µm). C. Hoescht (cyan) stained sagittal section of spinal cord from SCI animal with no scaffold. DA-traced axons (yellow) can be seen in the corticospinal tract (rostral aspect on the left, scale bar 500 µm). D. Hoescht (cyan) stained sagittal section of spinal cord from SCI animal implanted with uncoated lignocellulosic scaffold (outlined by white dotted line). DA-traced axons (yellow) can be seen in the corticospinal tract (rostral aspect on the left, scale bar 500 µm). E. Hoescht (cyan) stained sagittal section of spinal cord from SCI animal implanted with PLO-coated lignocellulosic scaffold (outlined by white dotted line). DA-traced axons (yellow) can be seen in the corticospinal tract (rostral aspect on the left, scale bar 500µm).

### Neural cell infiltration and axonal sprouting inside lignocellulosic biomaterials

Histological analysis of spinal cord tissue revealed host cell infiltration into the scaffolds from the rostral and caudal interface. Within the scaffold, both cell bodies and axonal projections expressing β-III tubulin, an early neuronal marker, were identified (Fig 5A, B). This finding suggests that endogenous adult neural stem cells infiltrated the biomaterial and initiated differentiation towards neural lineages. In addition, a substantial number of NF200-positive cells were observed inside the scaffold and surrounding tissue (Fig 5C-E). In both PLO-coated and uncoated scaffolds, NF200-positive cells densely clustered at the outermost edge of the scaffold and some NF200-positive neurites extended into its channels. To determine the effect of PLO coating on infiltration of NF200-positive cells, the distance of infiltration into the channels was measured from the scaffold edge towards the injury epicenter in NF200-stained sagittal sections. NF200-positive cells penetrated significantly further into the channels of PLO-coated scaffolds compared to those without the coating (p=0.0286, Fig 5F). On average, the longest distance of NF200-positive cell infiltration was 388.8±154.5µm into the PLO-coated scaffolds and 215.6±22.3µm in the uncoated scaffolds. However, the percentage of channels infiltrated by cells did not differ between the two conditions (p=0.8286). In PLO coated scaffolds, 93±0.8% of the channels were infiltrated by NF200-positive cells, on average, and uncoated scaffolds had 93±2% of their channels infiltrated. Interestingly, at the rostral and caudal interfaces of the biomaterial we observed a cluster of neurofilament-positive projections branching from the dorsal to ventral aspect of the spinal cord, perpendicular to the axis of the tracts (Fig 5D).

**Figure 5.**
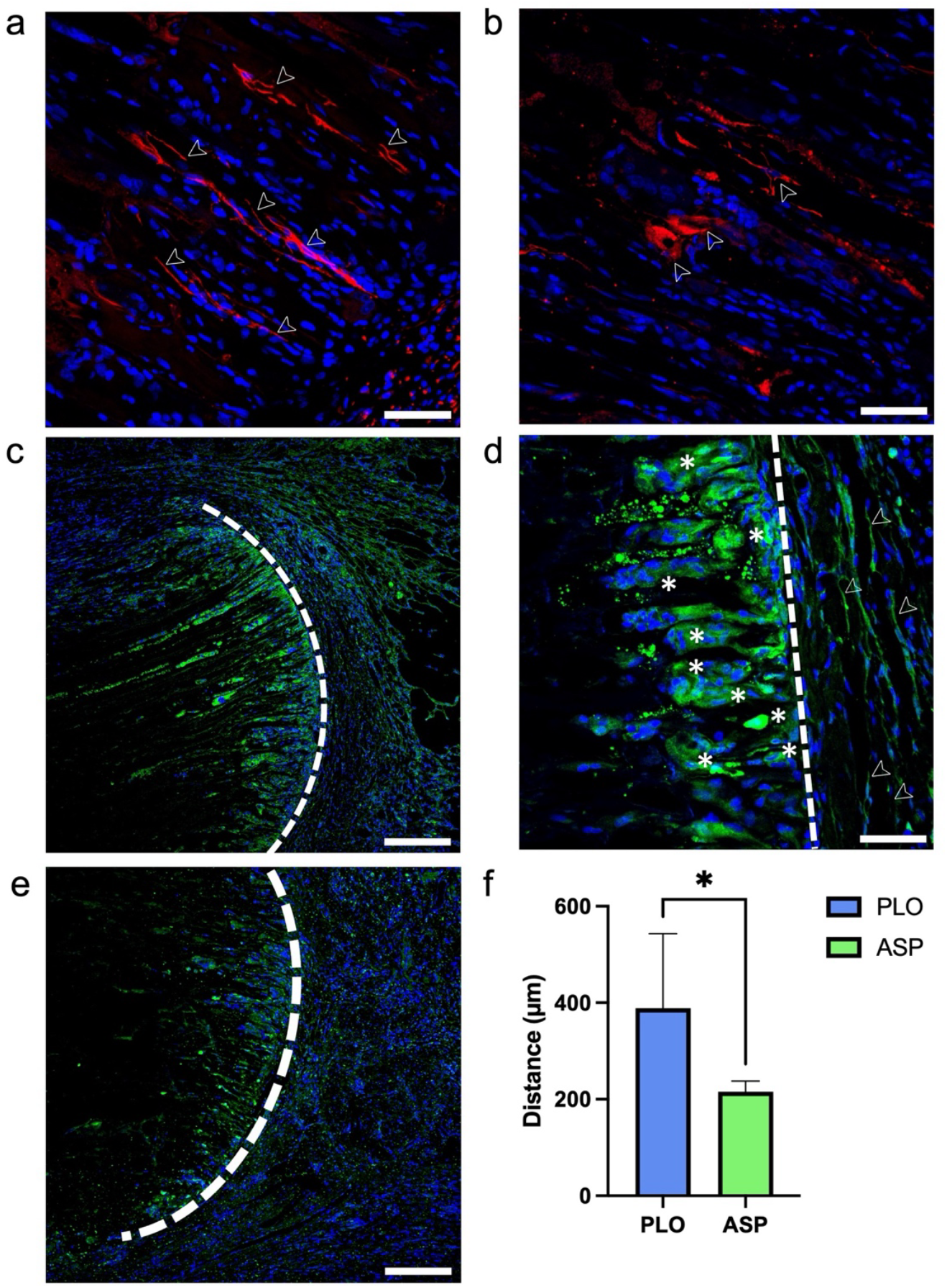
Immunostaining for ß-III tubulin and neurofilament-200 reveals neural cells attaching to scaffold and migrating along channels. A. ß-III tubulin (red) and Hoescht (blue) staining of sagittal section within PLO-coated biomaterial. Cell bodies (identified by arrow) can be seen inside the scaffold (scale bar 50µm). B. ß-III tubulin (red) and Hoescht (blue) staining of sagittal section within PLO-coated biomaterial. Axons (identified by arrow) can be seen sprouting inside the scaffold (scale bar 50µm). C. Neurofilament 200 (green) and Hoescht (blue) staining of sagittal section at the interface (dashed line) between PLO-coated biomaterial and spinal cord (scale bar 200µm). D. Neurofilament 200 (green) and Hoescht (blue) staining of sagittal section at the interface (dashed line) between PLO-coated biomaterial and spinal cord. NF200 positive cells can be seen infiltrating the biomaterial (asterisk) & NF200 axon projections extend from the dorsal to ventral aspect of the biomaterial (arrow) (scale bar 50µm). E. Neurofilament 200 (green) and Hoescht (blue) staining of sagittal section at the interface (dashed line) between uncoated biomaterial and spinal cord (scale bar 200µm). F. Longest distance of infiltration of NF200-cells into the scaffold for each experimental group. The mean ± SD of each group is shown (Unpaired T-test Mann Whitney; P=0.0286, n = 4 PLO animals, n = 4 ASP animals).

Finally, tissue sections from each experimental group were stained with luxol fast blue (LFB) to evaluate axon myelination in the injury site (Fig 6 and Supplementary Information Fig 6). As expected with a traumatic spinal cord injury, severe myelin loss was observed in the perilesional tissue, consistent with Wallerian degeneration. However, in animals that received a PLO-coated scaffold, dark blue bands of LFB staining were consistently observed at the rostral and caudal interface of the PLO-coated scaffold and tissue, indicating possible remyelination induced by the PLO. The intensity of the blue staining was quantified (methodological details and results described in the Supplementary Information Fig 6), and the results are consistent with the above observations. The average intensity of the blue signal for the PLO group was about two times greater than the No Tx group (p=0.002) and the untreated scaffold ASP group (p=0.012). Whereas the ASP and No Tx groups were not significantly different (p=0.417). It is important to note that uncertainty does arise in the quantification of these images due to the severely damaged nature of the tissue, animal variability and scaffold orientation during sectioning. Moreover, with the lack of a scaffold in the control group there is no direct comparison to the accumulation of signal around a scaffold. To address this uncertainty, we have included the entire image data set that was quantified (Supplementary Information Fig 5) to provide the reader with complete appreciation of what we are able to qualitatively observe by eye.

**Figure 6.**
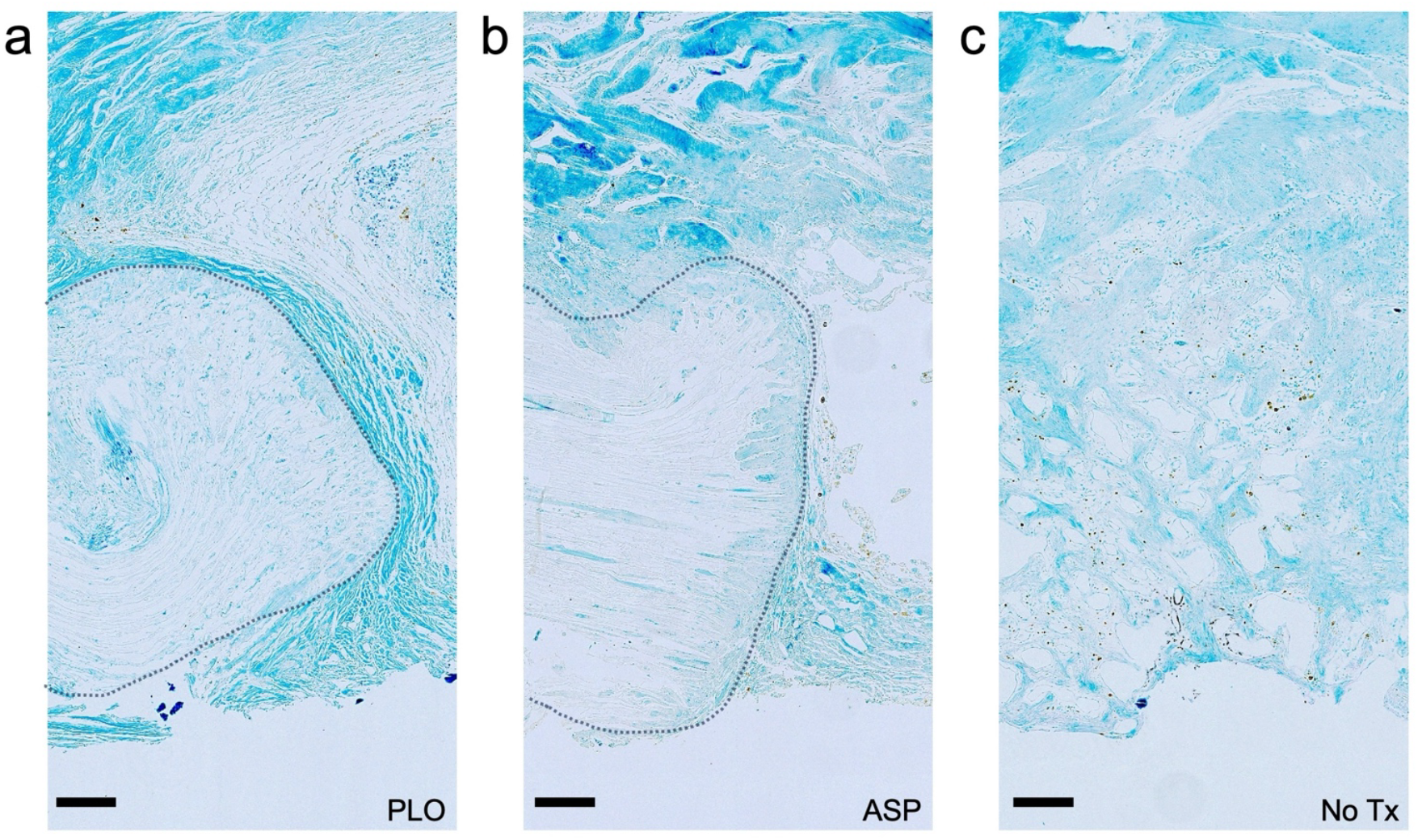
Luxol fast blue (LFB) staining of sagittal tissue sections for assessment of myelin at the site of spinal cord injury. Representative LFB staining at scaffold-tissue interface in animals implanted with PLO coated lignocellulosic scaffold. Outline denotes lignocellulosic scaffold (Scale bar = 200µm). B. LFB staining at scaffold-tissue interface in animals implanted with uncoated lignocellulosic scaffold (outlined) (Scale bar = 200µm). C. LFB staining at cyst-tissue interface in animals with no scaffold (Scale bar = 200µm).

## Discussion

Complete SCI leads to the loss of both sensorimotor and autonomic function distal to the injury site due to the interruption of ascending and descending pathways. Regrowth of axons and restoration of appropriate synaptic connectivity after SCI remains a major medical challenge. Various therapeutic strategies have attempted to induce sprouting of the corticospinal tract or to establish relay neural circuits that can mediate functional recovery^70,71^. Neural circuits in the injured spinal cord are known to exhibit plasticity after treadmill training^72^ or electrical epidural stimulation^73^ that can produce some degree of motor recovery. However, irregular synaptic connectivity after SCI can be maladaptive and lead to spasticity or neuropathic pain^74^. In recent decades, biomaterial-based therapeutic strategies for SCI have been explored to support axonal growth and deliver cells or pharmaceuticals locally to modulate the inhibitory milieu of the lesion. For instance, some groups have produced bioactive scaffolds that use cell signaling molecules to enhance axonal regrowth, myelination, and functional recovery^20^ while others have 3D printed scaffolds with microarchitectures that mimic the structures in the spinal cord to support the formation of neural relays^19^. As well, several scaffolds have been used as carriers to deliver various types of stem cells to the injured cord to promote repair^75^.

In the present study, we investigated the ability of naturally occurring microarchitectures found in plant-derived scaffolds to provide physical support to regenerating axons and guide their extension across a lesion in the spinal cord. Our results show that plant-derived scaffolds can become vascularized after implantation into the spinal cord and that endogenous neural cells expressing β-III tubulin and NF200 readily migrate into the channels of the scaffold in a linear orientation along the rostral-to-caudal axis. Aside from directing the extension of regenerating axons, the biomaterial may have contributed to motor recovery by its interaction with astrocytes, which are known to play a key role in SCI pathology. After SCI, activated microglia induce changes in the gene expression and morphology of astrocytes surrounding the lesion. The process of reactive astrogliosis is controlled by complex signaling pathways, notably NFκB or TGF-β signaling ^76^, and eventually leads to the formation of a dense glial scar. We found positive immunostaining for glial fibrillary acidic protein (GFAP) in sagittal spinal cord sections from each experimental group (Supplementary Information Fig 7). In all animals, GFAP expression was observed in the tissue rostral and caudal to the injury, with little to no GFAP visible in the injury epicenter. By occupying space in the injury site, the lignocellulosic scaffold may have reduced the number of reactive astrocytes migrating into the lesion and interfered with signaling mechanisms involved in the propagation of reactive astrogliosis, thereby attenuating tissue damage and enhancing motor ability. In addition, the presence of channels at the surface of the lignocellulosic scaffolds may have affected the alignment and morphology of astrocytes in the injury site, as previous reports have shown that cytoskeletal modulation in astrocytes can be induced by microgrooves^77^.

We tested the ability of plant-derived lignocellulosic scaffolds to support repair in combination with a coating of poly-L-ornithine (PLO), a positively charged synthetic amino acid that adheres to cellulose by electrostatic interactions. PLO is commonly used in neural stem cell culture and previous studies have demonstrated its ability to improve cell attachment and migration^78,79^. Specifically, PLO was shown to promote filopodia formation in neural progenitors in vitro^79^ by increasing the expression of α-Actinins 4, a critical effector of structural plasticity. Based on this, we hypothesized that PLO may have a similar effect in vivo after SCI and that increased sprouting of severed axons could assist in establishing neural relays across the scaffold, thereby mediating functional recovery.

We found that animals implanted with the uncoated scaffolds had a modest recovery of motor function in the BBB assessment and this recovery was enhanced with a PLO coating. By contrast, no significant recovery of motor function was observed amongst the No Tx animals during the recovery period. We utilized scaffolds with an average length of 2.9±0.5mm, which was customized to fit the specific dimensions of each animal’s injury. For the animals’ final BBB score, no dependence was observed on the length of scaffold (Supplementary Information Fig 8). For the animals’ retrograde tract tracing results, no dependence was observed on the length of scaffold (Supplementary Information Fig 9). Despite the motor improvements seen in biomaterial-treated animals, we did not find evidence of serotonergic activity below the injury. Immunohistochemistry staining for 5-HT revealed some sprouting of serotonergic axons, though positive 5-HT staining was limited to tissue on the rostral side of the complete transection in all experimental groups (Supplementary Information Fig 10). To investigate possible axon regeneration which may contribute to the observed motor recovery, we performed retrograde tract tracing by injecting CTb into the sciatic nerve. This neural tract tracing experiment demonstrated that sensory axons extended a greater distance into the injury site in animals with PLO-coated scaffolds compared to those without a scaffold. Our findings are consistent with those of Schackel et al, who found that PLO/laminin-coated hydrogels implanted in animals with a cervical hemisection had increased host cell migration and a slight increase in neurite growth^78^. Further experiments may elucidate the mechanism by which the PLO-coated scaffold produced this effect. It is unclear whether the CTb-traced sensory fibers in the injury site are regenerated axons or spared fibers that may result from reduced axonal retraction after injury. We speculate that the presence of the lignocellulosic scaffold in the injury site may slightly reduce cystic cavitation, thereby mitigating further axon retraction after injury. In addition, it is possible that the PLO coating promoted neurite extension, as demonstrated by other works^78–80^. Similarly, the immunostaining experiments we performed on sagittal sections of the scaffold after 12-weeks in vivo revealed NF200-positive cells migrating further into the channels of scaffolds with the PLO coating compared to the uncoated biomaterial. Taken together, our results highlight the potential of plant-derived scaffolds, though further work is necessary to optimize therapeutic strategies using these biomaterials with PLO or other peptides to further enhance neural cell infiltration to potentially enable communication between circuits below and above the injury.

The differences we identified in the sensory axons in PLO-treated animals could possibly have contributed to their recovery in motor ability, given the importance of sensory feedback in motor function after SCI^58,59,81,82^. To test whether these axons serve as the substrate for functional recovery, future experiments could be performed to assess the functionality of theses regenerated sensory fibers and to identify whether they have relevant synaptic contacts. In addition to sprouting of sensory axons, we observed what appeared to be enhanced myelination in the injury site of animals with PLO-coated scaffolds only. This was evidenced by positive LFB staining demonstrating myelin in the scaffold periphery which was not observed in the No Tx or ASP groups.

In a recent study, the ability of PLO to enhance myelin repair was demonstrated in an animal model of focal demyelination^83^. We speculate that myelin repair may be a contributing mechanism underlying PLO-induced motor recovery in our SCI model. Further, the positive effect of PLO on myelination in the injury site may be two-fold, as myelin breakdown products are known to be inhibitors of neuronal plasticity. Hence, by enhancing remyelination, PLO may also be creating a more permissive environment for axonal sprouting, consistent with our observations. In future experiments, plant lignocellulosic scaffolds could be seeded with stem cells, functionalized with neurotrophins or axon guidance molecules such as netrins or ephrins to further promote regeneration of axons through the scaffold.

Environmental enrichment (EE) was included as part of our multifaceted neurorehabilitation strategy due to its ability to enhance neuroplasticity and attenuate SCI-induced neuropathic pain ^84–86^. Environmental enrichment is a housing manipulation that includes novel toys, tunnels, nesting materials, puzzles, and running wheels along with opportunities for socialization with conspecifics. By enhancing plasticity, enriched environments support functional improvements in animal models of stroke and SCI^84–91^. Studies have indicated that EE promotes plasticity through various mechanisms including increased production of neurotrophic factors and changes in dendritic-spine density ^92^. In rodent SCI models, enrichment potentiates the regenerative ability of neurons via Creb-binding protein-mediated histone acetylation, which increases expression of regeneration-associated genes^90^. Many groups have reported that environmental enrichment substantially improves sensory and motor recovery after contusive SCI^84,86^. Similarly, in animal models of stroke, exposure to an enriched environment causes neuroanatomical changes including dendritic remodeling, axonal sprouting, and the release of growth factors ^93^. It has also been reported that environmental enrichment may promote white matter recovery after stroke by reducing microglia activation^94^. Altogether, environmental enrichment shows great promise as part of a multimodal SCI therapeutic strategy. In the present study, all animals were housed with EE and no standard housing (SH) controls were included, since SH is an impoverished environment that is not representative of the conditions of human SCI patients. In addition to improving animal welfare throughout the study, we speculate that the enriched environment may have contributed to the growth of axons after SCI by expanding the regenerative ability of sensory neurons, as previously reported^85^. Enrichment has been explored as a therapeutic modality for stroke and traumatic brain injury, with results suggesting it may act synergistically with other treatments to enhance functional outcomes^95^. However, it is possible that enrichment in our study overshadowed the benefits of the PLO-coated scaffold alone.

Importantly, due to the architecture of the scaffolds utilized in this study, their mechanical properties are highly anisotropic. The elastic modulus of the scaffold is about 10-fold softer when measured perpendicular to the long axis (n=10), compared to measurements made parallel (n=10) to the long axis (12±4 kPa vs 148±53kPa). At a high level, one can imagine the scaffold to behave similarly to a bundle of straws which will bend easily when deformed perpendicular to the long axis. However, the same bundle of straws will be able to withstand higer forces when compressed parallel to their long axis. Reported values for the elastic modulus of spinal cord tissue can vary by orders of magnitude, with some reporting up to 40 kPa for human spinal cord tissue^48^. There is significant variation in the literature regarding the elastic modulus, depending on the specific anatomical region being tested, the species, the testing method and sample preparation^48–51^. Factors such as the test temperature and removal vs non-removal of the dura mater greatly affect the overall mechanical characteristics of the tissue. Therefore, it is difficult to confidently point to one specific range to elastic moduli that characterize a “healthy”” spinal cord. Despite this, materials being investigated for SCI repair also report elastic moduli in a broad range, even as high as 260– 300 kPa^19^. However, it is still important in tissue engineering to strive to match the physical properties of the tissues under investigation with any implanted biomaterials. The scaffolds utilized in this study do broadly fall within the range of reported values for the spinal cord. A key consideration is that in most studies the “bulk modulus” is typically being reported. As our previous studies have shown^34,36,38,41,96^, the bulk modulus is typically significantly higher than the local modulus (as measured with atomic force microscopy). It is precisely this local nanoscale elastic modulus, which is what cells are actually “feeling” and responding to. Regardless of the considerations above, in the work presented here, and in our previous studies^37^, the scaffolds we have utilized are clearly able to support neuronal growth and proliferation, modest motor recovery, tissue integration and tissue regeneration. Therefore, the evidence thus far suggests that while the mechanical properties of these scaffolds may be further optimized in the future, at present they do not appear to prevent some degree of recovery.

Biodegradability can also be important in biomaterial design; however, it must also be carefully controlled to ensure that it aligns with the regeneration rate of axons, does not result in a loss of tissue integrity, and the process does not create toxic byproducts. While other non-degradable biomaterials have been explored for neural tissue engineering applications^97^, they are sometimes associated with challenges such as chronic inflammation, gliosis and scarring. Prior to completing the present study, we performed a pilot study in which uncoated ASP scaffolds were implanted in a cohort of animals for up to 28 weeks. In pilot, PLO coated scaffolds were not tested and the animals did not receive enrichment which makes the data difficult to directly compare to the current study. However, the pilot study does enable the investigation of potential chronic inflammatory immune responses and adverse events caused by the long-term implantation of a non-degradable scaffold (Supplementary Information Fig 11). Importantly, after 28-weeks, the implants were well integrated into the spinal cord and there was no evidence of chronic inflammation, granuloma or rejection. While there is a concern that plant-derived scaffolds are largely non-degradable, we speculate that it is precisely due to their inertness that they can offer advantages in some tissue engineering applications. The scaffold provides cells and tissues with a highly stable platform on which to regenerate. This can be particularly important in cases of severe trauma in which the tissue microenvironment is highly damaged and inflammatory. Moreover, as these scaffolds are carbohydrate based, if degradation does occur one of the main by-products would be glucose.

Taken together, it is clear in this first proof-of-concept study that plant-derived scaffolds can potentially have an important role to play in the treatment of SCI. While, future studies should investigate opportunities to systematically understand the impact of utilizing softer/stiffer scaffolds, or the introduction of scaffolds that are able to resorb, this study provides a starting foundation and motivation on which to further expand this body of work. Importantly, this study has also demonstrated the potential of functionalizing plant-derived scaffolds with PLO as part of a multimodal SCI therapeutic strategy which could also include environmental enrichment. We found that coating the biomaterial with PLO supported hindlimb motor recovery and neural tissue repair in a rat model of complete transection. The biomaterial was infiltrated by endogenous neural cells, which migrated along the linearly oriented channels of the plant scaffold. Retrograde neural tracing highlighted regeneration of sensory tracts in the spinal cord after treatment with PLO-coated scaffolds. Overall, our results point to potential future treatment strategies that may one day utilize plant-derived scaffolds in combination with other therapeutics and physical interventions.

## Methods

### Biomaterial production

Asparagus (Asparagus officinalis) was purchased from local supermarkets (Loblaws, Ottawa). The asparagus was stored at 4°C in the dark for a maximum of one week and kept hydrated. To prepare the lignocellulosic scaffolds, the asparagus (with a diameter 14–17 mm) were washed, and the end of the stalks was cut to remove any dried tissue. A 4 mm biopsy punch was used to cut out cylindrical sections close to the edges of the tissue to maximize the number of vascular bundles in each scaffold. Effort was made to avoid the central fibrous tissues common in all angiosperm plants. The resulting lignocellulosic scaffolds were 4mm in diameter and were cut to various lengths (1mm, 1.5mm, 2mm, 2.5mm, 3mm, 3.5mm and 4mm in length) using a microtome blade (Westin Instruments Boston) and verified with a Vernier caliper. The scaffolds were then placed into a 50ml Falcon tube containing 0.1% sodium dodecyl sulphate (SDS) (Sigma-Aldrich). Samples were shaken for 72 hours at 180 RPM at room temperature. Scaffolds were then transferred into new sterile individual microcentrifuge tubes, washed and incubated for 12 hours in phosphate-buffered saline (PBS). Following the PBS washing steps, the asparagus were then incubated in 100 mM CaCl_2_ for 24 hours at room temperature and washed 3 times with dH2O. Samples were then sterilized in 70% ethanol overnight. Finally, scaffolds were washed 12 times in a sterile saline solution. In some cases, scaffolds were also incubated overnight in a 100µg/ml solution of poly-L-ornithine (30-70kDa, Sigma-Aldrich, P3655).

### Young’s Modulus Testing

Scaffolds were loaded onto a CellScale UniVert (CellScale) compression device. The Young’s modulus was measured by compressing the material to a maximum of 10% strain at a compression speed of 50 µm/s. The force-indentation curves were converted to stress-strain curves and the Young’s modulus was extracted from the elastic region of the curves.

### Animal Housing and Environmental Enrichment

All procedures described in this study were approved and performed in accordance with standards set out by the University of Ottawa Animal Care and Veterinary Services ethical review committee (Approved Protocol Number SCe-3701). Methods and results are reported in accordance with ARRIVE guidelines. Juvenile female Sprague Dawley rats were purchased from Charles River. Rat tickling was performed with the juvenile rats as a habituation technique to improve welfare. Experimenters followed the Panksepp method^98^ of rat tickling which consists of 15 sec rest followed by 15 sec of dorsal contacts and pins for a total of 2 minutes for 4 days of training. Upon completing tickle training, every rat was tickled before every procedure. Clicker training was performed with the juvenile rats to encourage cooperation in all behavioral assays and to develop positive affect with experimenters. Briefly, rats were encouraged to step onto a plastic platform and given food rewards upon successful completion, accompanied by a ‘click’. Clicker training was done for 5 days (4 mins daily). All rats were housed in pairs throughout the study and got 20 minutes of daily group play with conspecifics. During group play, up to 10 rats were placed in a 1-meter diameter arena with climbing structures, tunnels, foraging toys, food enrichment and nesting material. Enrichment was provided in all rats’ home cages, including a wooden block, nylon bone (Bio-Serv, K3580), bunny block (Bio-Serv, F05274) and metal swing. In addition to a diet of standard rat chow, all rats were given daily food enrichment including mini yogurt drops (Cedarlane Bio-Serve, F7577), banana chips (Cedarlane Bio-Serve F7161), fruity bites (Cedarlane Bio-Serve F6038), ABC fruit blend (Cedarlane Bio-Serve F7228), Mealworms (Cedarlane Bio-Serve 9264), veggie-bites (Cedarlane Bio-Serve F5158) and pumpkin (E.D. Smith).

### Spinal Cord Transection Surgical Procedure

Once rats reach a weight of 250-300g, they are anesthetized with isoflurane USP-PPC and injected subcutaneously with normal saline (Baxter) and enrofloxacin (Baytril). Laminectomies were performed at the T8-T9 level to expose the spinal cord, followed by a dorsal midline durotomy. The entire cord was gently lifted by a hook before cutting with micro scissors. Surgifoam 1972 (Ethicon) was used to establish hemostatic control, then the gap was measured after 10 minutes to select the appropriately sized lignocellulosic implant. We utilized scaffolds with a diameter of 4 mm and an average length of 2.9±0.5mm which was customized to fit the specific dimensions of each animal’s injury. Prior to surgery, animals were be split into four groups: sham (laminectomy only, n=2), No Tx (full transection only, no scaffold, n=8), ASP treated (lignocellulosic implant only, n=9), and PLO treated (PLO-coated lignocellulosic implant, n= 11). PLO-coated scaffolds were rinsed twice with sterile water before implantation. While implanting the scaffold, ARTISS fibrin sealant (Baxter) was applied into the cavity. The muscle and adipose tissue were reapproximated with 3-0 Vicryl sutures (Johnson & Johnson) and the skin was closed with Michel clips (Fine Science Tools). Rats received post-operative care including bladder expressions 4 times daily, pain monitoring and management with buprenorphine HCl, when necessary, weight loss and dehydration tracking.

### Biocompatibility assessment by subcutaneous implantation

Sprague Dawley rats, aged 6-8 weeks (n=12), received pre-operative analgesia via a subcutaneous injection of buprenorphine (0.05 mg/kg) and were anesthetized using 2% Isoflurane USP-PPC. The animals were placed on a heating pad, and a large area of hair was shaved off from their dorsal aspect. The skin was scrubbed with 4% chlorhexidine gluconate followed by Soluprep (2% chlorohexidine gluconate and 70% ethanol). Four 1 cm-long incisions were made lateral to the midline. A scaffold was implanted into each of the four subcutaneous spaces and the incisions were sutured closed using 6-0 Prolene. Bupivacaine 2% (transdermal) was applied to each incision and rats were transferred to a recovery incubator at 35°C with oxygen. Buprenorphine SR 1mg/kg was administered 4 hours after recovery and enrofloxacin (Baytril) 10ml/kg was administered for 3 days perioperatively. Scaffolds were resected at 4-, 8- and 12-weeks post implantation, after euthanasia by CO2. The implants were resected including a 2 cm perimeter of healthy skin and transferred into 10% formaldehyde (Sigma-Aldrich) for 48 hours. Samples were then transferred to 70% ethanol and stored at 4°C. Histological analysis was performed on paraffin embedded tissue. Serial sections were cut (5µm thick) and stained with either hematoxylin-eosin (H&E) or Masson’s trichrome (MT). Micrographs were captured using Zeiss MIRAX MIDI Slide scanner equipped with a 40X objective. Images were analyzed by a pathologist to assess cell infiltration, extracellular matrix deposition, and vascularization.

### BBB Locomotor Assessment

Functional recovery of the hindlimbs was assessed by a weekly BBB open field assessment.^99^ Each rat was placed in a 1-meter diameter arena covered with a non-slippery floor and recorded by 5 cameras. The 4-minute videos were then scored by three blinded observers and the average of three examiners was calculated for each animal at each timepoint. Spasticity and movement occurring simultaneously with urination was ignored and confirmed with repeated views of the videos. Two weeks after receiving the spinal cord injury, one animal was excluded based on their BBB score, which was above 5.

### KSAT Swim Assessment

Before SCI, all rats were acclimatized to the pool daily (up to 5 minutes) for 5 days. During this pre-training, each rat was placed into a clear acrylic swim tank (depth 20cm, length 150cm) filled with water (27-30°C) and encouraged to swim via clicker training. Animals with poor performance were excluded from the study before the SCI surgery. After rats sustained a SCI, the swim assessment was performed every 2 weeks to assess functional recovery. Each animal was allowed three runs across the swim tank with 20 second rests between runs. Swimming was recorded by 4 cameras and videos were scored by 3 blind observers using the Karolinska Institutet Swim Assessment Tool (KSAT).^62^

### Inclined Plane Assessment

Before the spinal cord injury, all animals were trained to perform the inclined plane test as described by Rivlin et al^*63*^. Each rat was placed on an inclined plane while adjusting the slope to determine the maximum angle at which the animal can maintain its position without falling. Every 2 weeks after the spinal cord injury, the inclined plane test was performed to examine sensorimotor recovery.

### Retrograde Tract Tracing of Ascending Sensory Afferents

Animals also received 2uL of Cholera Toxin Subunit B conjugated to Alexa Fluor 647 (CTb, ThermoFisher, cat. C34778, 1% solution in sterile PBS) injected bilaterally into their sciatic nerves (4uL total per rat). Rats were anesthetized with 3% isoflurane, a linear incision was made along the femur, and the gluteus maximus was separated from the gluteus medius to expose the sciatic nerve. Using a surgical microscope, a nick was made in the proximal portion of the exposed nerve. The CTb solution was loaded into a Hamilton syringe, and the needle was inserted 5mm into the sciatic nerve. CTb was injected in increments of 0.5µL, waiting 30 seconds and withdrawing the needle by 1mm per injection. Once the injection was complete, a sterile Q-tip was used to remove any CTb that was expelled from the nerve and the incision was sutured using 4-0 Prolene sutures. Post-operatively, 2% bupivacaine was applied topically and animals were given buprenorphine (0.05mg/kg subcutaneously) and Gabapentin (Chiron, 50mg/kg subcutaneously) for analgesia. Pain was assessed TID and additional buprenorphine was administered as necessary. To prevent infections, Enrofloxacin (Bayer, 10mg/kg subcutaneously) was administered daily for 3 days peri-operatively.

### Anterograde Labeling of the Corticospinal Tract

11 weeks post-operatively, rats from each group were injected with neural tracers to visualize axons in and around the injury site. Animals were anesthetized with 3% isoflurane and placed into a stereotactic frame. After a surgical scrub, a sagittal incision was made to expose the top of the skull and the periosteum was scraped off. Using a Zeiss surgical microscope, the skull was levelled by measuring the Dorsal-Ventral axis at 4 random points and ensuring they are within 0.05mm of each other. Bregma was located and its coordinates were used to calculate the target location of each injection. Using a surgical drill (Micro Drill, Harvard Apparatus) attached to the stereotactic frame, 8 holes were drilled into the cranium. Stereotactic injections were made into the right and left motor cortex at coordinates: Injection 1: Anterior-posterior (AP) −0.5mm, Medial-lateral (ML) ±2mm, Dorsal-Ventral (DV) 1.5mm. Injection 2: Anterior-posterior (AP) −1mm, Medial-lateral (ML) ±2.5mm, Dorsal-Ventral (DV) 1.5mm. Injection 3: Anterior-posterior (AP) −1.5mm, Medial-lateral (ML) ±2mm, Dorsal-Ventral (DV) 1.5mm. Injection 4: Anterior-posterior (AP) −2mm, Medial-lateral (ML) ±2.5mm, Dorsal-Ventral (DV) 1.5mm. Injections were made using a 5uL Hamilton syringe. After a 2-minute delay, a volume of 0.5µL of 10% Dextran Amine Alexa Fluor 488 (ThermoFisher, cat. D22910, 10,000 MW) was injected into each location at a rate of 250nl/min for a total injection volume of 4µL per rat. Each injection was followed by a 2-minute delay to ensure diffusion into the tissue. Once all 8 injections were complete, the scalp was sutured with 4-0 Prolene sutures and 2% transdermal bupivacaine was applied to the incision. Animals had 2 weeks of recovery time before euthanasia to allow for transport of neural tracers. For tissue collection, animals were deeply anesthetized with isoflurane USP-PPC and euthanized by cardiac perfusion with 500ml of 1xPBS followed by 500ml of 4% paraformaldehyde. The brain and spinal cord were dissected out and fixed overnight in 4% paraformaldehyde at 4°C, then stored in 70% ethanol at 4°C until embedding and sectioning.

### Quantification of neuroanatomical tract tracing

Cross-sections of spinal cord tissue above and below the injury site were imaged by confocal laser scanning microscopy to confirm successful tracer uptake. For retrograde tracing, CTb-traced axons were seen in the dorsal columns of T10 cross-sections. For anterograde tracing, dextran amine labelled axons were identified within the corticospinal tract of T6 cross-sections. For each animal, 3 sagittal sections in the middle of the cord were imaged by confocal laser scanning microscopy and the distance between the tracers and the injury epicenter was measured using ImageJ (Supplementary Information Fig 12). For retrograde tracing with CTb (n= 4 PLO animals, n=2 ASP animals, n=3 No Tx animals), the distance (mm) was measured between the furthest rostral CTb-traced axon and the injury epicenter. For anterograde tracing with dextran amine (n=4 PLO animals, n=4 ASP animals, n=3 No Tx animals), the distance (mm) was measured between the furthest caudal dextran amine-traced axon and the injury epicenter. In animals implanted with a scaffold, the injury epicenter was deemed to be at the mid-point of the scaffold (½ the total length of the scaffold). In ‘no scaffold’ control animals, the injury epicenter was deemed to be at the mid-point of the syrinx at the lesion site (½ the total length of the cyst). Averages were calculated for each group and a one-way ANOVA was performed.

### GFAP Immunostaining

Paraformaldehyde-fixed paraffin-embedded tissue sections (5µm thick) were deparaffinized and pre-treated using heat-mediated antigen retrieval with Sodium Citrate buffer (pH 6.0). Slides were then rehydrated in 1X TBST buffer and blocked for 30 minutes with Rodent Block R (Biocare RBR962H). Sections were then incubated with Rabbit GFAP (1:3000, Sigma AB5804) at room temperature for 1.5 hours. Sections were washed with 1XTBST and then incubated with the Goat anti-Rabbit-488 antibody for 2 hours in the dark at room temperature. This was followed by incubation with a quencher (Vector TrueView Autofluorescence Quenching Kit #SP-8400, Vector Labs) to decrease autofluorescence. Sections were then washed, incubated with 5 ug/ml of DAPI (ThermoScientific #62248) and coverslipped.

### β-III Tubulin Immunostaining

Paraformaldehyde-fixed paraffin-embedded tissue sections (5µm thick) were deparaffinized and tissue was permeabilized at room temperature (5 mins) the following buffer: 0.5% Triton-X, 20mM HEPES, 300mM Sucrose, 50mM NaCl, 3mM MgCl2, 0.05% sodium azide. Sections were incubated in blocking buffer (6% Normal Goat Serum in 1xPBS) for 10 minutes. Sections were rinsed twice in 1xPBS before incubation at 4°C overnight in mouse anti-β-III tubulin antibody (10ug/ml, MAB1195, R&D systems). The following day, after 2 washes of 1xPBS, sections were incubated at room temperature (2.5 hours) in goat anti-mouse alexa Fluor 594 polyclonal antibody (1:200, A11005, Thermofisher) and counterstained with Hoescht (1:2000).

### Neurofilament 200 Immunostaining

Paraformaldehyde-fixed paraffin-embedded tissue sections (5µm thick) were deparaffinized and tissue was permeabilized at room temperature (3x 5 minutes) in 1X TBST. Sections were incubated in blocking buffer (5% Normal Goat Serum in 1X TBST) for 30 minutes followed by 3 washes in 1X TBST. Tissue was incubated at 4°C overnight in rabbit anti-NF200 (1:3000 dilution in 1X PBS, cat. N4142 Sigma). The following day, after 2 washes of 1X PBS, sections were incubated at room temperature (2 hours) in goat anti-rabbit alexa fluor 488 (cat. A11008 Thermofisher) and counterstained with Hoescht (1:2000).

### 5-HT Immunohistochemistry

Paraformaldehyde-fixed paraffin-embedded tissue sections (5µm thick) were deparaffinized and incubated in rabbit anti-serotonin antibody (1:2500 dilution in 1X PBS, Sigma S5545) for 2 hours. Immunohistochemistry was performed using DAB as the chromogen and counterstained with Hoescht (1:2000).

### Luxol Fast Blue Staining

Paraformaldehyde-fixed paraffin-embedded tissue sections (5µm thick) were deparaffinized and hydrated to 95% alcohol. Staining was done in a 0.1% LFB solution, overnight at 37°C. Sections were rinsed in 95% alcohol followed by distilled water. Sections were then differentiated by a brief dip in a 0.05% lithium carbonate solution, then in 70% alcohol followed by 3 rinses in distilled water.

### Statistical Analysis

Experimental data were analyzed using GraphPad Prism. For the distance of infiltration of NF200-positive cells, the differences between two groups were compared utilizing the Student’s t-test at a significance level of P < 0.05 & the Mann Whitney test was performed. Differences between multiple groups were compared by one way ANOVA followed by Tukey’s multiple comparisons test (for neuroanatomical tract tracing) or the uncorrected Fisher’s LSD (for BBB locomotor assessment) at a significance level of P < 0.05.

## Acknowledgements

The authors would like to thank the Animal Care and Veterinary Service staff and the University of Ottawa Louise Pelletier Histology Core (RRID: SCR_021737) for their continuous support in this project. We would like to thank Dr. Rafay Azhar for advising on histopathological interpretation. This work was supported by the Li Ka Shing Foundation (AEP), the Natural Sciences and Engineering Research Council (TB and AEP), Canada Research Chairs Program (AEP), the Canada Foundation for Innovation (AEP), the Canadian Institutes of Health Research (TB), the Canada Graduate Scholarship Program (LJC) and the Ontario Graduate Scholarship Program (LJC).

## Author contributions statement

LJC, KLAW, DJM, CMC, ECT, TB and AEP conceived the experiment(s). LJC, KLAW, AB, DJM, CMC, MLL, RJH, RB and RM conducted the experiment(s), LJC, KLAW, DJM, CMC, MLL, RJH, R-JKO, IS, AG, TB and AEP authors analysed the results. All authors reviewed the manuscript.

## Data Availability

The datasets used and/or analysed during the current study available from the corresponding author on reasonable request.

