## Supplementary Information for "Poly-L-Ornithine Coated Plant Scaffolds Support Motor Recovery in Rats after Traumatic Spinal Cord Injury"

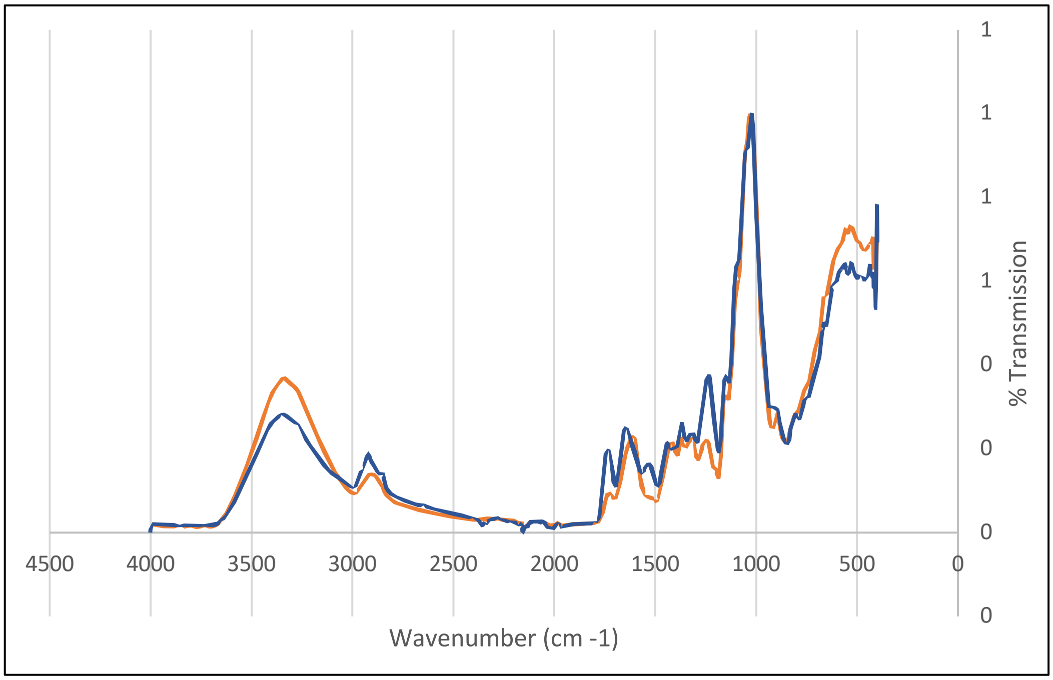


**Supplementary Figure 1.** Fourier-transform infrared spectroscopy (FTIR) spectra of uncoated scaffolds (orange) and PLO-coated scaffolds (blue). Samples were frozen at -80°C overnight and lyophilized for 24h before being analyzed using the KBr disk method. Spectra were recorded using a Nicolet 6700 AT-FTIR from 4000 to 500 cm-1.

A substantial body of research has focused on characterizing cellulosic materials using various analytical methods, with FTIR spectroscopy being one of the most common techniques (Hinterstoisser et al., 2000; Kokot et al., 2002; Guo & Wu, 2008). FTIR offers valuable insights into the supramolecular architecture of lignocellulosic scaffolds and enables the determination of chemical compositions in both native and modified cellulose, hemicellulose, and lignin fractions. However, one complexity is that these carbohydrate polymers share similar chemical moieties resulting in many overlapping vibrations modes that will be present in the FTIR spectra.

With that said, the scaffolds utilized in this study are a lignocellulosic material, which is constituted of cellulose, hemicellulose and lignin. Among these three major biopolymers, cellulose serves as the principal reinforcing component of the plant cell wall. It is organized into microfibrils, where extensive hydrogen bonding between cellulose chains results in a robust and highly ordered crystalline structure. Hemicellulose, a heterogeneous polysaccharide, surrounds the cellulose microfibrils and contributes to matrix flexibility and crosslinking, while lignin provides rigidity and hydrophobicity, enhancing structural integrity and resistance to microbial degradation. It should be noted that poly-l-ornithine shares many characteristic vibrational modes associated with C-H, CH2, N-H and C=O that will overlap with characteristic regions of lignocellulosic materials. However, some insights can still be gained from FTIR analysis. The peaks between 3000 cm-1 and 3800 cm-1 are attributed to hydrogen bonding between hydrogen and oxygen. The most complex region of cellulose FTIR spectra is the fingerprint region from 1430 cm-1 to approximately 850 cm-1, which contains signals from the numerous sp3 single bond vibrational modes. These observations are highly consistent with studies of other lignocellulosic materials (Kostryukov et al., 2023; Li et al., 2018).

**Supplementary Table 1:** Assignment of transmittance bands to characteristic polymer bonds.

| Wavenumber ($cm^{-1}$) | Assignment | Polymer |
|---|---|---|
| 3600 to 3000 | Hydrogen-bonded O-H stretching.<br><br>N-H stretching | Cellulose, hemicellulose, lignin.<br><br>PLO |
| 3000 to 2800 | C-H asymmetrical and symmetrical stretching.<br><br>Distinct peaks for alkanes and presence of CH2 groups | Cellulose, lignin, pectin.<br><br>PLO side chain |
| 1750-1700 | C=O stretching | Lignin and PLO |
| 1650-1550 | O-H bending (absorbed water) | Cellulose |
| 1500 to 1200 | Hydrocarbon groups (CH2 scissoring)<br><br>C-N stretching and bending | Cellulose, hemicellulose, lignin, PLO<br><br>PLO |
| 1100-1000 | C-O-C stretching at β-glycosidic linkage<br><br>C-C, C-OH, C-H ring and side group vibrations | Cellulose |
| 895 | COC, CCO and CCH deformation and stretching | Cellulose |

a

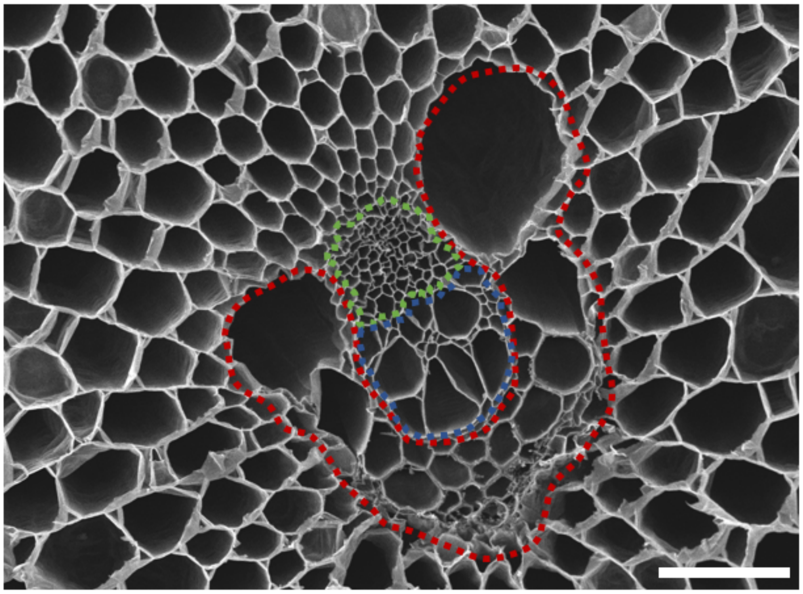


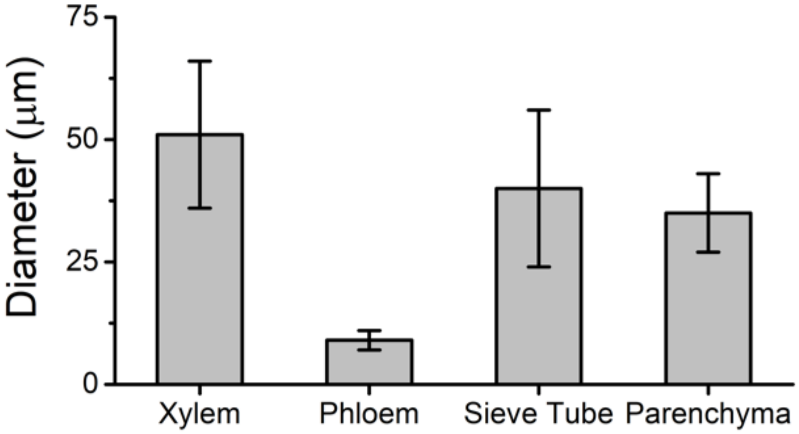


**Supplementary Figure 2. Different structures of the vascular bundle. a)** A scanning electron microscope image of the surface of a decellularized scaffold revealing a single vascular bundle (VB) and surrounding parenchyma tissue (Scale bar = 100μm). The elements of the VB are highlighted to show the distinct channel architectures. The xylem (red) are channels that run the entire length of the asparagus and transport water within the plant. The phloem (blue) transport sugars within the asparagus from photosynthetic cells to non-photosynthetic cells. Phloem structures differ from xylem as they contain highly perforated sieve elements along their length. The sieve tubes are outlined in blue. The sieve tubes contain specialized cells with no nucleus that have roles in transporting carbohydrates/ messaging molecules throughout the plant. **b)** Characteristic diameters of the various elements of the vascular bundle, xylem channels (51±15μm), sieve tubes (40±16μm), parenchyma (35±8μm) and the phloem (9±2μm).

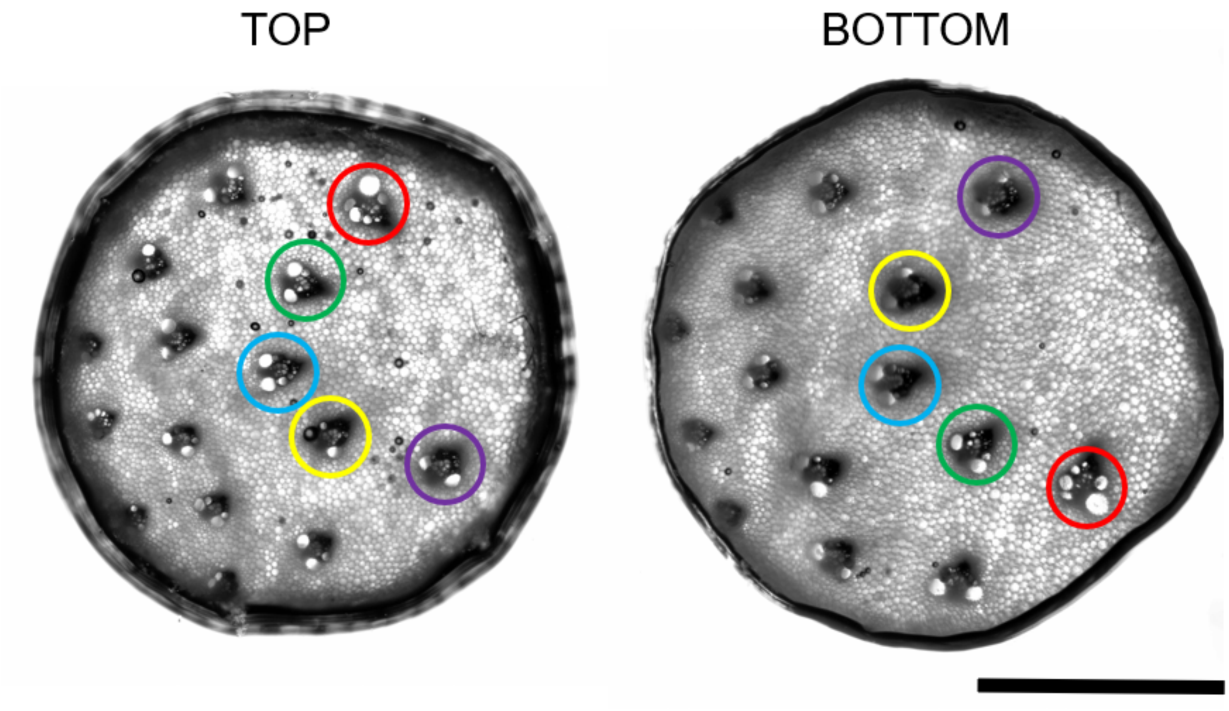


**Supplementary Figure 3. The two opposite ends of the scaffold.** Phase contrast image of the entire surface of the scaffolds revealing the distribution of the VB within the scaffold on the top surface and the emergence of the same VB in the nearly the exact position in the bottom surface (Scale bar = 2mm).

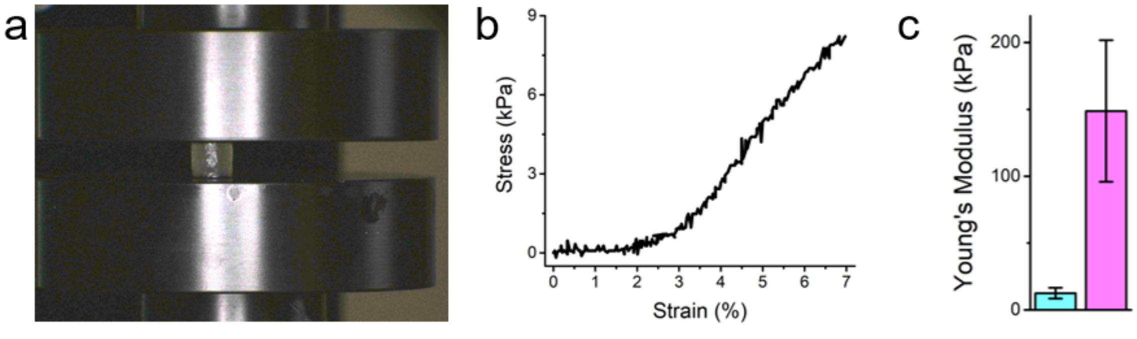


**Supplemental Figure 4. The young modulus of the SCI scaffold prior to implantation. a)** The SCI scaffold loaded into a CellScale UniVert compression platform. **b)** The elastic deformation of the stress strain curve fort the SCI scaffold used to determine the scaffold. **c)** The quantified young's modulus of the SCI scaffold along the perpendicular axis (blue, n=10) and parallel to the long axis (pink, n=10).

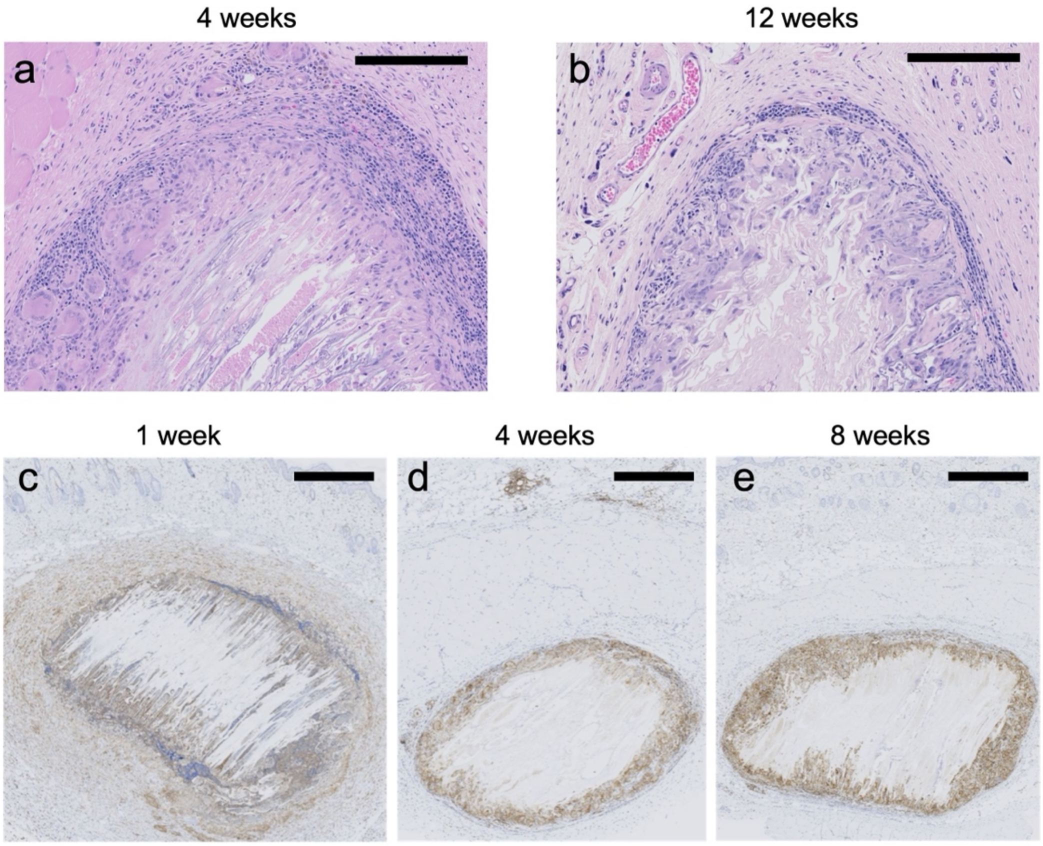


**Supplementary Figure 5. In vivo biocompatibility of PLO-coated cellulose implant.** A. Hematoxylin & eosin staining of sagittal section of subcutaneous implant at 4 weeks post-implantation (Scale bar = 250µm). B. Hematoxylin & eosin staining of sagittal section of subcutaneous implant at 12 weeks post-implantation (Scale bar = 250µm). C. CD45 immunohistochemistry staining of sagittal sections showing hematopoietic cells (in brown) infiltrating the biomaterial 1 week post-implantation (Scale bar = 900µm). D. CD45 immunohistochemistry staining of sagittal sections showing hematopoietic cells (in brown) infiltrating the biomaterial 4 weeks post-implantation (Scale bar = 900µm). E. CD45 immunohistochemistry staining of sagittal sections showing hematopoietic cells (in brown) infiltrating the biomaterial 8 weeks post-implantation (Scale bar = 900µm).

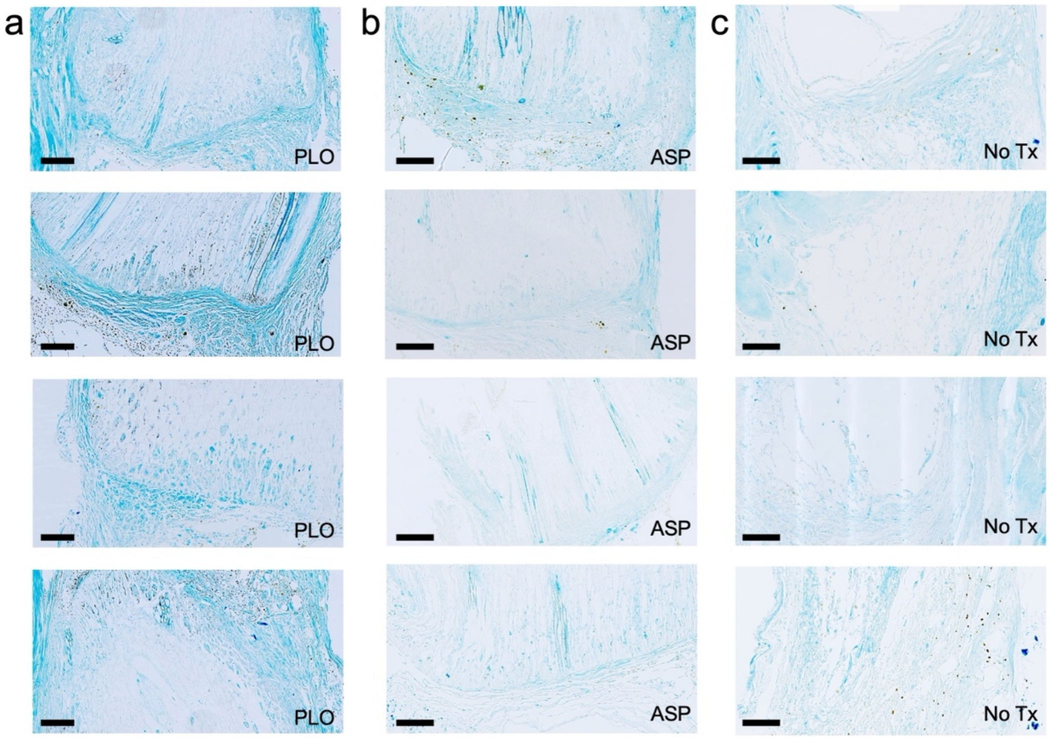


**Supplementary Figure 6. Luxol fast blue (LFB) staining of sagittal tissue sections for assessment of myelin at the site of spinal cord injury in an additional 12 animals.** a) Representative LFB staining at scaffold-tissue interface in n=4 animals implanted with PLO coated cellulose scaffold. Outline denotes cellulose scaffold (Scale bar = 200µm). b) LFB staining at scaffold-tissue interface in n=4 animals implanted with uncoated cellulose scaffold (outlined) (Scale bar = 200µm). c) LFB staining at cyst-tissue interface in n=4 animals with no scaffold (Scale bar = 200µm).

Qualitatively, regions of more intense LFB staining can be observed around the PLO scaffolds, with more diffuse signal in the untreated scaffold and control groups. While considerable variability can occur as it is difficult to section in the exact same plane and orientation from animal to animal, it is tempting to quantify the data. Added to the complexity is the lack of any scaffold in the control groups leading to images of a severely traumatized tissue with little regeneration observed. With the above caveats in mind, RGB (red-green-blue color model) images were first converted to HSB (hue-saturation-brightness color model) in ImageJ. The saturation channel was then extracted and a constant threshold applied to isolate regions of blue signal. The resulting binary image was utilized to create a mask, and the mask applied to an inverted greyscale version of the original RGB image to extract the intensity of the blue signal in those regions. On a scale from 0 to 255, the average intensity of the blue signal for the control group was 38.90 ± 3.99, the untreated scaffold group was 49.03 ±24.74 and the PLO group was 91.97 ± 9.85. A one-way ANOVA with post-hoc Tukey test confirms that the PLO group possessed significantly more blue intensity than the untreated scaffold ($p=0.012$) and the control ($p=0.002$). Furthermore, the untreated scaffold and control groups were not significantly different ($p=0.417$).

Though we hypothesize that the PLO may enhance motor recovery by promoting remyelination, some confounding factors may be contributing to the recovery observed in this study. For example, the environmental enrichment provided in the animals' cages could produce functional improvements by enhancing neuroplasticity or increasing expression of regeneration-associated genes (Berrocal et al., 2007; Clemenson et al., 2015; Hutson et al., 2019; Johansson & Belichenko, 2002; Koopmans et al., 2012; Mering & Jolkkonen, 2015; Neves et al., 2023; Zhu et al., 2021). It is also possible that spontaneous motor recovery resulted from sprouting of spared fibers or intraspinal reorganization. The scaffold or PLO coating may have influenced the immune response and led to sparing of tissue below the injury or otherwise influenced plasticity in lumbar spinal cord circuits, leading to improved motor function.

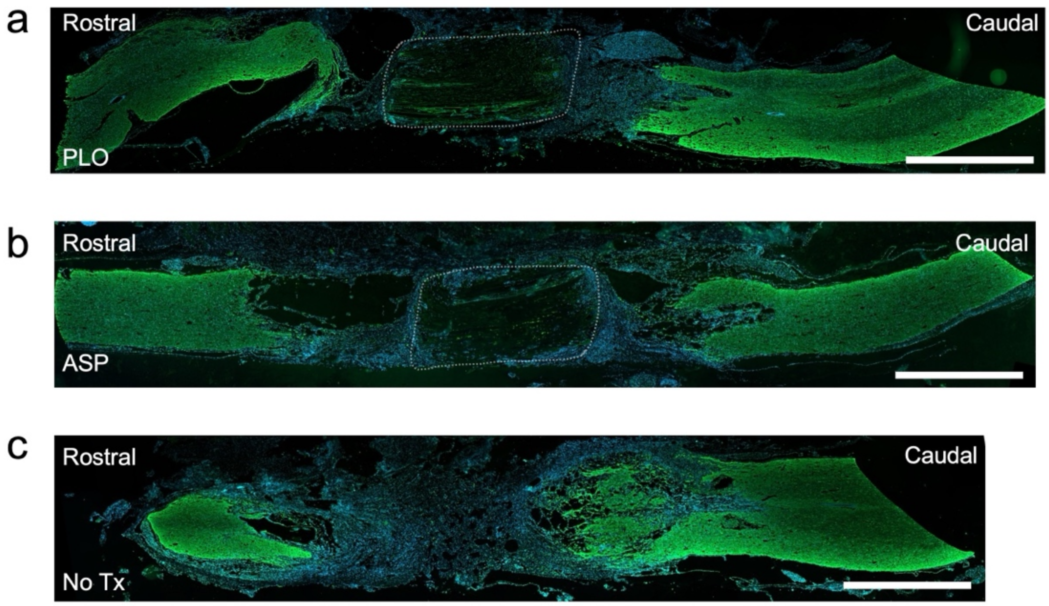


**Supplementary Figure 7. Immunostaining for glial fibrillary acidic protein (GFAP) in sagittal spinal cord sections after SCI.** a) GFAP (green) stained cord section from animal implanted with PLO coated cellulose scaffold. Nuclei stained with dapi (blue). Outline denotes cellulose scaffold (Scale bar = 200µm). b) GFAP (green) stained cord section from animal implanted with uncoated cellulose scaffold (outlined). Nuclei stained with dapi (blue). (Scale bar = 200µm). c) GFAP (green) stained cord section from animal with no scaffold. Nuclei stained with dapi (blue) (Scale bar = 200µm).

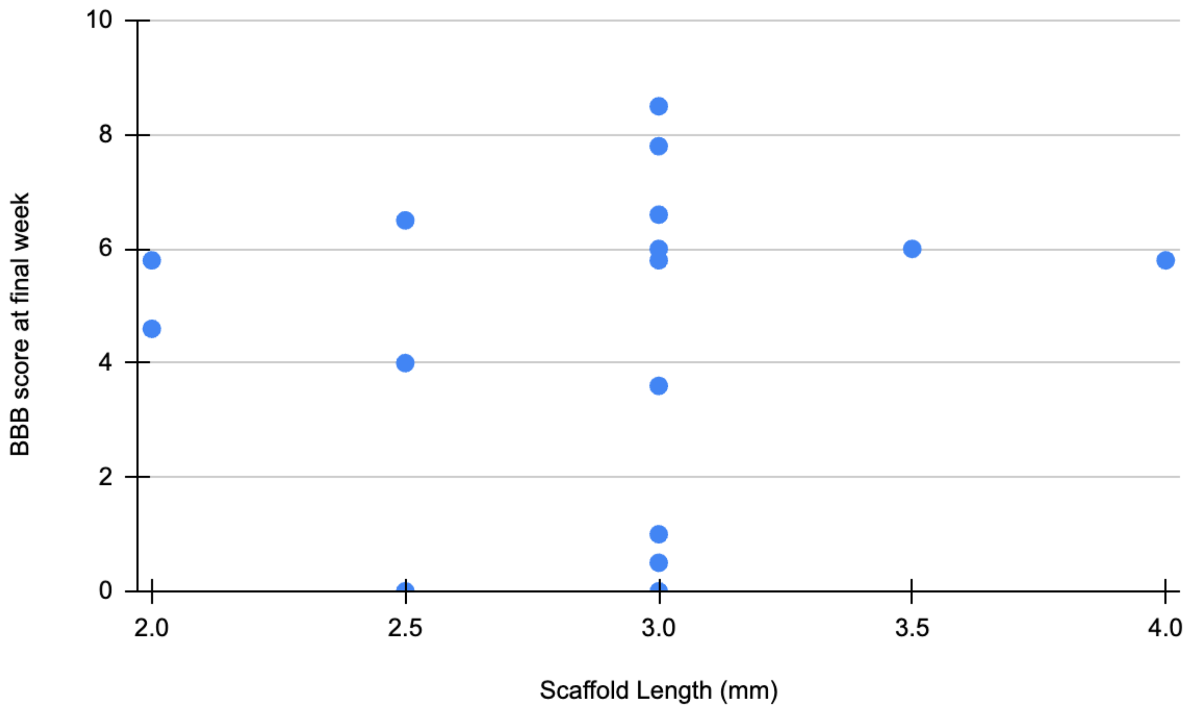


**Supplementary Figure 8. Scatter plot of relationship between scaffold length and final week BBB score.** Each point represents one animal (n=17). Scaffold lengths were measured at time of implantation. Average length of scaffolds was 2.9±0.5mm, which was customized to fit the specific dimensions of each animal's injury.

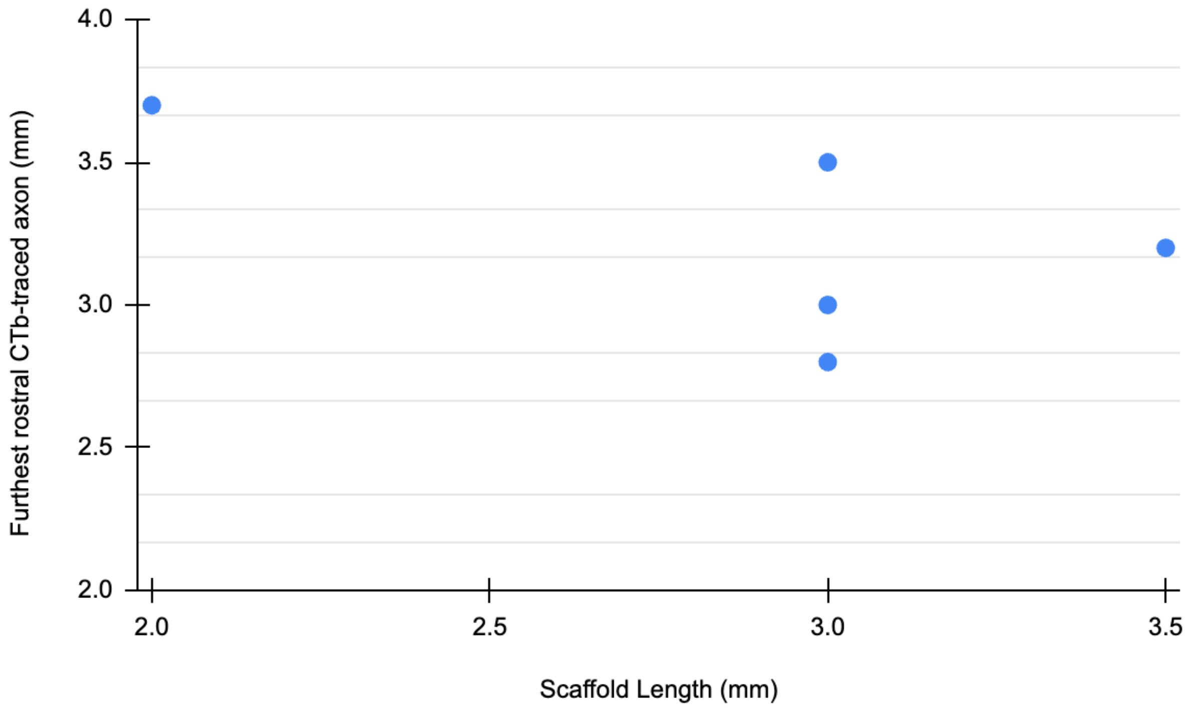


**Supplementary Figure 9. Scatter plot of relationship between scaffold length and retrograde tract tracing results (Furthest rostral CTb-traced axon (mm)).** Each point represents one animal (n=5). Scaffold lengths were measured at time of implantation. Average length of scaffolds was 2.9±0.5mm, which was customized to fit the specific dimensions of each animal's injury.

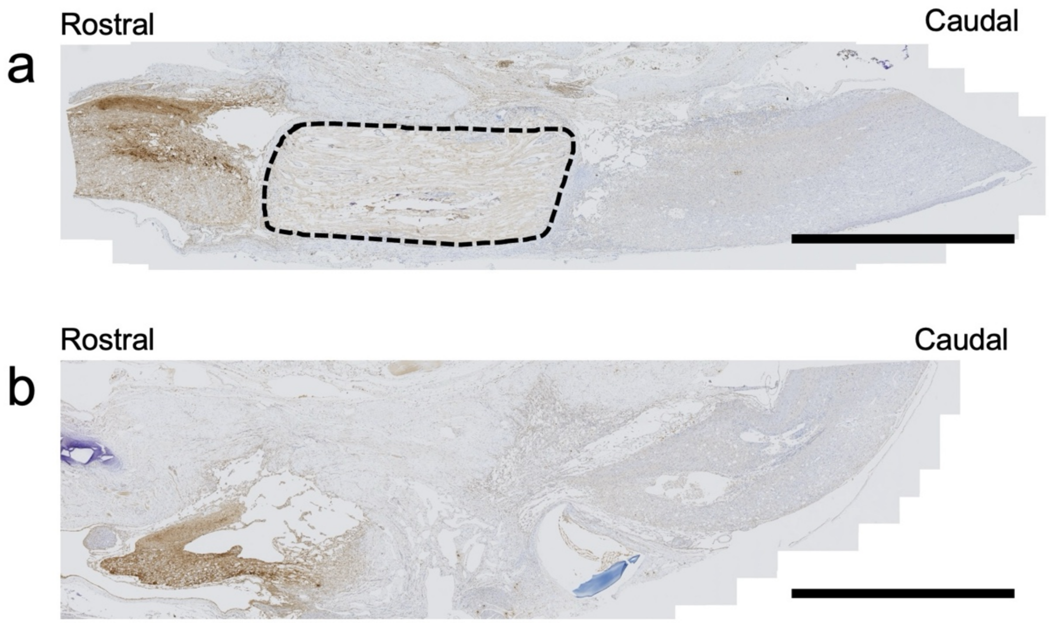


**Supplementary Figure 10. 5-HT immunohistochemistry staining of sagittal section of T8-T9 spinal cord injury.** a) 5-HT staining of spinal cord implanted with ASP scaffold (outlined). IHC was performed using DAB as the chromogen (brown) (Scale bar = 2mm). b) 5-HT staining of spinal cord with no implant. IHC was performed using DAB as the chromogen (brown) (Scale bar = 2mm).

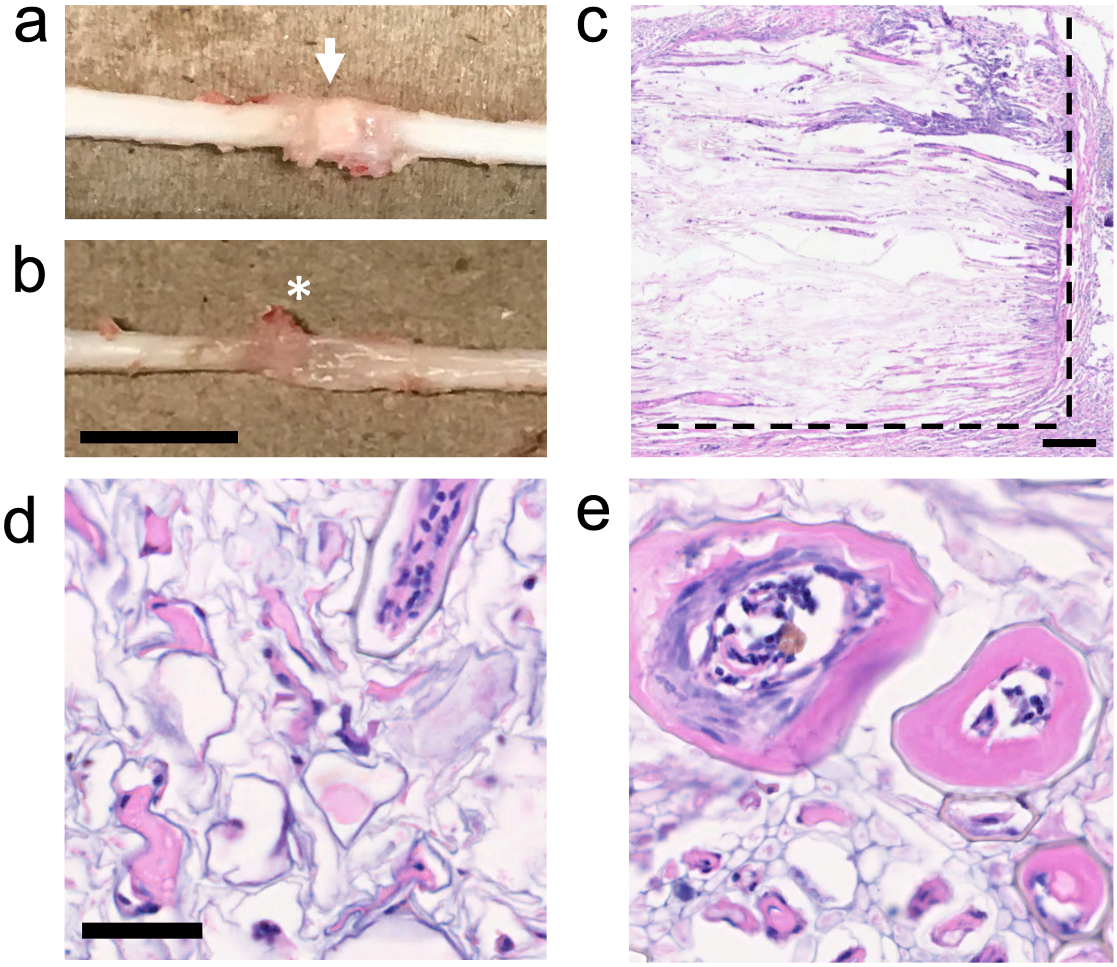


**Supplementary Figure 11. Scaffold implantation at 28-weeks.** A much earlier pilot study utilizing uncoated scaffold was performed prior to the current study. Although the methods were similar, PLO was not investigated, nor was animal enrichment provided making the results difficult to directly compare to the current study. However, the study does provide insight on the impact of long-term implantation of a non-degradable plant-derived scaffold. In this case, n=11 animals had an uncoated scaffold implanted into the legion created by a full transection. After 28 weeks, some degree of motor recovery was observed, consistent with the current study. **a)** More importantly, during dissection, the scaffold (arrow) was observed to be well integrated and could not be removed from the two stumps of the spinal cord. The scaffold was so well adhered that it could support the weight of the entire CNS when lifted. **b)** The control groups were difficult to remove as the stumps loosely adhered via scar and connective tissues (asterisk). The damage to the surrounding spinal cord tissue can also be observed much further in both the rostral and caudal spinal cord stumps (Scale bar = 1cm and applies to both). **c)** Sagittal view of hematoxylin-eosin staining of a scaffold at the T8-T9 vertebrate after 28 weeks of implantation. The vascular bundles can be seen throughout the scaffold, and many are infiltrated with host cells (Scale bar = 200 µm). Even after 28 weeks there is no significant degradation of the scaffold, and the borders of the scaffold are still apparent (represented in dashed lines). Representative high-resolution hematoxylin-eosin stained regions of the scaffold **d)** parenchyma and **e)** vascular bundle channels (Scale bar = 50 µm and applies to both) are presented. The images reveal the intact micro-architecture of the scaffold, nuclei of infiltrating cells and blood vessels with thick endothelial linings in the parenchyma tissue and vascular bundle. Overall, there was no evidence of chronic inflammation, rejection or any adverse events, highly consistent with several implantation studies our group has published in the past.

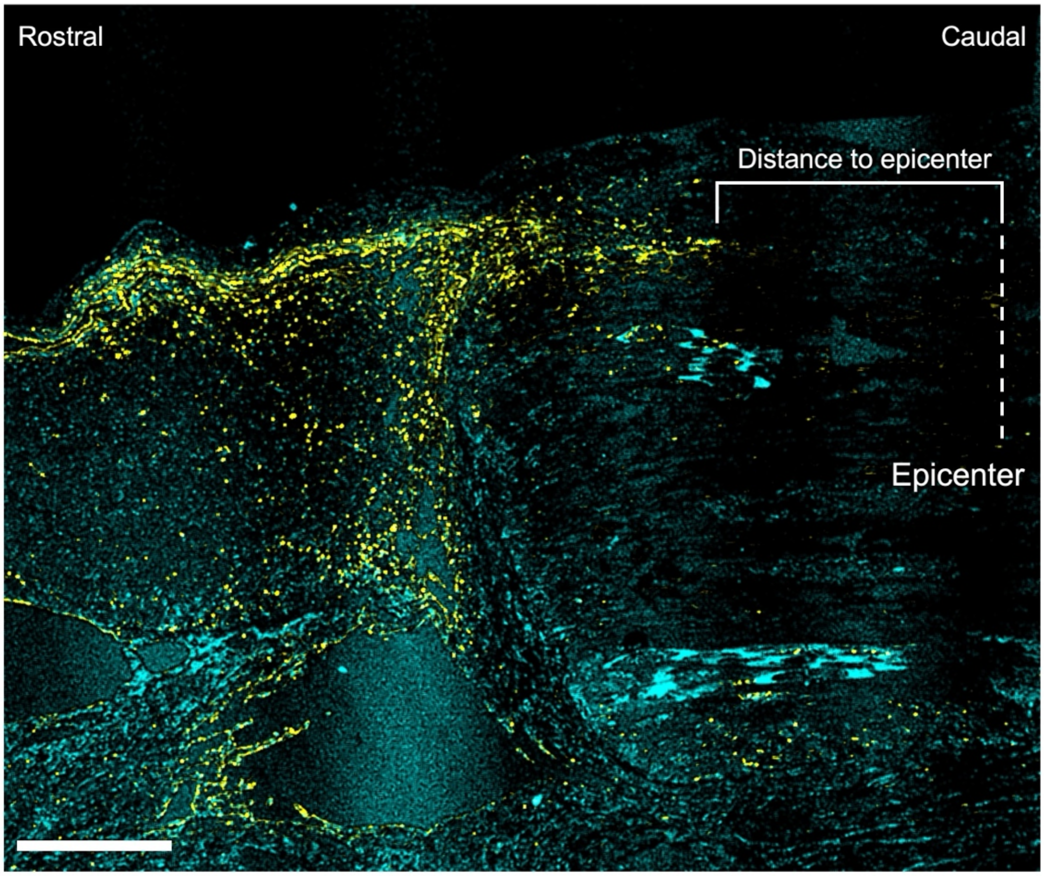


**Supplementary Figure 12. Tracer quantification in sagittal sections of T8-T9 spinal cord tissue.** For anterograde tracing with dextran amine (yellow), the distance (mm) was measured between the furthest caudal dextran amine-traced axon and the injury epicenter (indicated by dotted line). In animals implanted with a scaffold, the injury epicenter was deemed to be at the mid-point of the scaffold (half the total length of the scaffold). In 'no scaffold' control animals, the injury epicenter was deemed to be at the mid-point of the syrinx at the lesion site (half the total length of the cyst). All measurements were taken in ImageJ (Scale bar = 500 µm, nuclei in cyan).
